# Mono-ADP-ribosylation by ARTD10 restricts Chikungunya virus replication by interfering with the proteolytic activity of nsP2

**DOI:** 10.1101/2020.01.07.896977

**Authors:** Sarah Krieg, Fabian Pott, Laura Eckei, Maud Verheirstraeten, Mareike Bütepage, Barbara Lippok, Christine Goffinet, Bernhard Lüscher, Patricia Verheugd

**Affiliations:** Institute of Biochemistry and Molecular Biology, RWTH Aachen University, 52072 Aachen, Germany; Institute of Virology, Campus Charité Mitte, Charité – Universitätsmedizin Berlin, 10117 Berlin, Germany; Berlin Institute of Health, 10117 Berlin, Germany

## Abstract

A subset of intracellular mono-ADP-ribosyltransferases diphtheria toxin-like (ARTDs, aka mono-PARPs) is induced by type I interferons. Some of these mono-ARTDs feature antiviral activity while certain RNA viruses, including Chikungunya virus (CHIKV), encode mono-ADP-ribosylhydrolases, suggesting a role for mono-ADP-ribosylation (MARylation) in host-virus conflicts. CHIKV expresses four non-structural proteins (nsP1-nsP4), with nsP3 containing a macrodomain that hydrolyzes and thereby reverses protein MARylation *in vitro* and in cells. This de-MARylation activity is essential as hydrolase inactivating mutations result in replication defective virus. However, the substrates of MARylation during CHIKV infection are unknown and thus it is unclear how the macrodomain contributes to virus replication and how mono-ARTD-dependent MARylation confers antiviral immunity. We identified ARTD10 and ARTD12 as restriction factors for CHIKV replication in a catalytic activity-dependent manner. CHIKV replication requires processing of the non-structural polyprotein nsP1-4 by the nsP2-encoded protease and the assembly of the four individual nsPs into a functional replication complex. Expression of ARTD10 and ARTD12 resulted in a reduction of processed nsPs. Similarly, MAR hydrolase inactive CHIKV replicon mutants revealed a decrease in processed nsPs, comparable to an nsP2 protease defective mutant. This suggested that the macrodomain contributes to nsP2 protease activity. In support, a hydrolase-deficient virus was complemented by a protease-deficient virus. We hypothesized that MARylation regulates the proteolytic function of nsP2. Indeed, we found that nsP2 is MARylated by ARTD10. This inhibited nsP2 protease activity, thereby preventing polyprotein processing and consequently virus replication. This inhibition was antagonized by the MAR hydrolase activity of nsP3. Together, our findings provide a mechanistic explanation for the need of the viral MAR hydrolase for efficient replication of CHIKV.

**Author Summary:** Infectious diseases still pose major health threats. Especially fast evolving viruses find ever new strategies to manipulate the immune response. With climate warming and increased human mobility vector-borne pathogens like Chikungunya virus (CHIKV) spread and cause world-wide epidemics. Beyond the acute phase, CHIKV patients regularly suffer from chronic rheumatism. This entails a decline in life quality and an economic burden. To date no drugs are approved and the mode of pathogenesis remains elusive. Here we describe a mechanistic function of the CHIKV nsP3 macrodomain. We found that the viral nsP2 is mono-ADP-ribosylated interfering with its auto-proteolytic function. The nsP3 macrodomain removes this modification and restores the protease activity that is essential for replication. Because macrodomains are highly conserved they might represent broad antiviral targets.

## Introduction

Upon viral infection host cells initiate an antiviral immune response. Because the regulation of protein function through post-translational modifications (PTMs) is among the quickest mechanisms to adapt to challenges such as viral infections, PTMs display an essential part of antiviral signaling. Viruses have developed multifaceted strategies to evade or even hijack cellular mechanisms, e.g. encoding proteins that regulate PTMs and thereby counteract the antiviral reaction of the host [1–5]. Recent findings have elaborated on the role of intracellular ADP-ribosylation at the host-pathogen interface [6–12]. ADP-ribosylation is an ancient PTM of proteins and intracellularly mainly catalyzed by members of the ADP-ribosyltransferase diphtheria toxin-like (ARTD) family (also known as poly-ADP-ribosylpolymerases (PARPs)), which is composed of seventeen proteins [1, 13]. These enzymes use NAD^+^ as co-factor to transfer ADP-ribose onto a substrate protein with release of nicotinamide. Based on their catalytic features the ARTDs are subdivided into three classes. The first class contains ARTD1, 2, 5, and 6, which are capable of transferring multiple ADP-ribose moieties in an iterative process and thereby forming long polymers. This results in poly-ADP-ribosylation (PARylation) of substrate proteins. The second and largest class is defined by members that are restricted to mono-ADP-ribosylation (MARylation) (ARTD3, 4, 7-12, 14-17) [13–17], whereas the third group is defined by ARTD13, which lacks the ability to bind NAD^+^ and thus is catalytically inactive [18].

PARylation, arguably best known for its function in DNA damage response, has also been linked to chromatin organization, ribosome biogenesis, telomere maintenance, signaling processes and cell death [1, 19, 20]. In contrast functions of MARylation are less well understood but expanding with links to DNA damage repair, gene expression, signaling, stress response and cell death [1, 21]. Recent findings indicate a role for MARylation in host-pathogen conflicts [1, 21, 22]. The expression of several mono-ARTDs is triggered by type I interferons (IFNs) or pathogen-associated molecular patterns (PAMPs), like LPS, as part of an innate immune response to pathogens [6, 8, 11, 23–25]. Among these mono-ARTDs are ARTD7, ARTD10 and ARTD12, which have been identified to mediate clearance of Venezuelan Equine Encephalitis Virus (VEEV) by interfering with viral replication [7, 8]. In line with this, Zika virus replication is restricted by ARTD12 [26]. In addition to these molecular and cell-based studies, evolutionary analysis suggests a role for several ARTD family members at the host-pathogen interface, among them the macrodomain containing ARTDs (ARTD7-9) and ARTD13 [27]. Although catalytically inactive, the latter is best studied for being restrictive for viral replication by recognizing foreign CG-rich RNA through its Zinc finger domains [28, 29].

ADP-ribosylation can be propagated and regulated by macrodomains, structurally highly conserved protein folds found among all domains of life and a subset of positive single-strand RNA ((+)ssRNA) viruses [30, 31]. Generally, these macrodomains can function as readers or erasers of ADP-ribosylation. Whereas PAR chains are recognized and bound by the macrodomain of histone macroH2A1.1, the macrodomains 2 and 3 of murine Artd8 specifically interact with MARylated proteins [32, 33]. Degradation of PAR chains is mediated by PARG, which contains a macrodomain that cleaves the bond between single ADP-ribose units, but leaves the protein proximal ADP-ribose unaffected [34, 35]. ADP-ribosylation has been demonstrated to be a fully reversible modification with the identification of MacroD1, MacroD2 and TARG1 as hydrolases capable of removing MAR [36–38]. Besides these cellular MAR hydrolases, viral macrodomains have recently been characterized as MAR hydrolases. Macrodomains found in a subset of (+)ssRNA viruses, including members of the alphavirus genus such as Chikungunya virus (CHIKV), remove MARylation [6, 9, 10, 39]. This provides additional support for a role of MARylation in host-pathogen conflicts.

CHIKV is vector-borne and caused spreading epidemic outbreaks in Asia, Africa, the Americas and Europe until to date [40]. In addition to an acute flu-like phase that is associated with fever, rash and arthralgia, roughly a third of patients develop chronic joint rheumatism that can last up to several years [41]. Hence, this virus poses a growing threat with considerable life quality and economic burdens, especially because to date no vaccine or therapeutic has been approved, although first clinical vaccine trials are in progress [40, 42]. Therefore, it is crucial to further elucidate the function of the non-structural proteins to identify viral or cellular therapeutic targets for the CHIKV containment and treatment.

CHIKV encodes a non-structural polyprotein (nsP) that is translated early after infection. This protein is then cleaved into the 4 individual nsP1-4 that together form a functional replication complex. The processing occurs auto-catalytically through the protease that is part of nsP2 [43]. Mutations interfering with protease activity result in defective CHIKV replication [44]. Similarly, mutations interfering with the hydrolase activity, which is part of nsP3, abolish replication [9]. However, in contrast to nsP2, the biological function of the nsP3 macrodomain remains elusive.

Here we identified the interferon inducible mono-ARTDs, ARTD10 and ARTD12, as host factors restricting CHIKV replication. ARTD10 and ARTD12, dependent on their catalytic activity, prevent polyprotein processing. Similarly, the lack of a hydrolytically active macrodomain results in defective polyprotein processing. We identified nsP2 as substrate for MARylation *in vitro* and in cells. MARylation of nsP2 by ARTD10 inhibits its proteolytic activity, which supports our observation of defective polyprotein processing. We demonstrate that the MAR hydrolase activity associated with nsP3 removes the modification from nsP2, thereby reactivating its proteolytic activity. Together, our data provide a mechanistic explanation for the role of the CHIKV MAR hydrolase activity of nsP3 for viral replication and offers one mechanism of how MARylation functions in host-virus conflicts.

## Results

### ARTD10 and ARTD12 are restrictive for CHIKV replication

CHIKV relies on a functional macrodomain for replication [45]. This suggests that the capability to reverse MARylation is essential for proper virus replication and that mono-ARTDs function as antiviral host factors. To identify mono-ARTDs that potentially affect CHIKV replication we performed knockdown experiments (Fig. 1a). HEK293 cells were transfected with siRNA oligo pools targeting IFNα-inducible mono-ARTDs (Supplementary Fig. 1a and [6]), such as *ARTD10, ARTD12, ARTD8,* or *ARTD7*, prior to transfection with CHIKV replicon RNA. This replicon encodes the four non-structural proteins but lacks the open reading frame of the structural proteins. Instead, the subgenomic promotor of the replicon controls the expression of Gaussia luciferase, which we analyzed as surrogate for virus replication (Fig. 1b) [46]. Replication was analyzed at 9, 24 and 30 hours post transfection (hpt) (Fig. 1a). At the early time point *ARTD10* and *ARTD12* knockdowns showed a significant increase in replication, while the effect decreased at later time points. Knockdowns of *ARTD8* or *ARTD7* only allowed for minor increases in replication compared to control (Fig. 1a). Based on these findings we decided to focus on ARTD10 and ARTD12 and their role in CHIKV replication. We employed HEK293 cells stably expressing TAP-tagged ARTD10 or ARTD12, either wildtype (wt) or catalytically inactive mutants (Supplementary Fig 1b,c and [14]), to test whether overexpression of these proteins interferes with CHIKV replication (Fig. 1c-g). Protein expression was induced by doxycycline (Dox) treatment prior to transfection with replicon RNA. Doxycycline treatment itself had little effect on CHIKV replication (Supplementary Fig. 1d), still control cells were always treated with doxycycline as well. ARTD10 and ARTD12 reduced replication two-fold, while the respective catalytically inactive variants, ARTD10-G888W(GW) or ARTD12-H564Y(HY), showed little effect (Fig. 1c,d), suggesting that the MARylation activity of these enzymes is required to interfere with CHIKV replication. These differences are not attributed to variations in protein levels, as the mutants were expressed more efficiently compared to the wt proteins (Fig. 1e, Supplementary Fig. 1c). Similarly, HEK293 cells transiently expressing ARTD10 or ARTD12 showed reduced replicon replication, whereas no influence of the PARylating ARTD1 or ARTD14 enzymes, which do not respond to IFNα, was observed (Supplementary Fig. 1e,f). Further analysis revealed that ARTD10 and ARTD12 had additive repressing effects on replication when analyzed 30 hpt (Fig. 1f).

**Figure 1:**
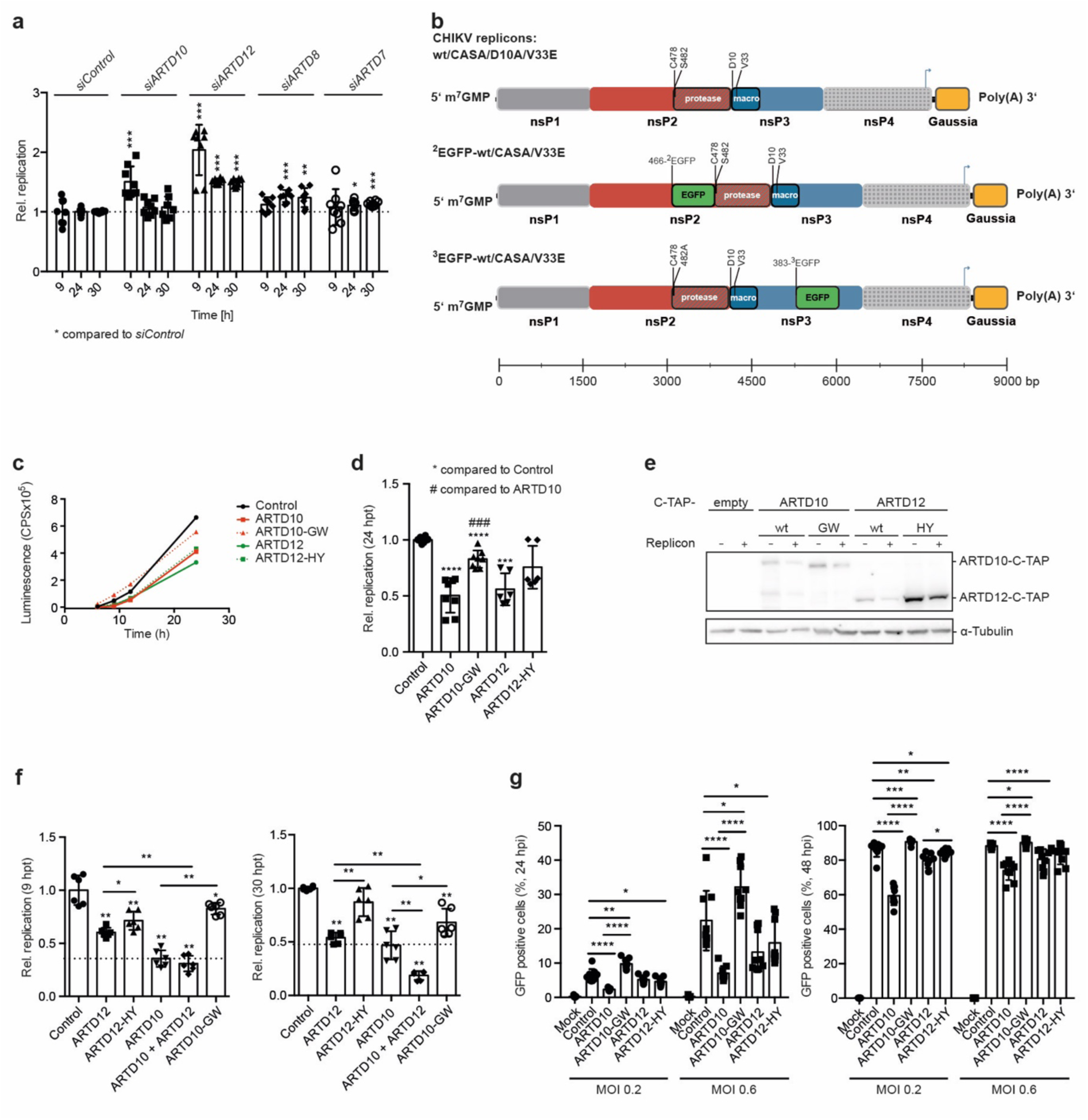
CHIKV replication is inhibited by ARTD10 and ARTD12 dependent on catalytic activity. *(**a**)* HEK293 cells were co-transfected with siRNA pools targeting the different *ARTD* mRNAs as indicated and with ^3^EGFP replicon RNA. Gaussia luciferase was analyzed 30 hpt. Normalization was against *siControl*. Error bars indicate SD (n = 4; 2 technical replicates measured per n; two-tailed Mann-Whitney test). *(**b**)* Schematic representation of the replicons used in this study. The scale bar indicates the length of the replicon variants in base pairs (bp). *(**c-e**)* HEK293 Flp-In T-REx cells stably expressing TAP-tagged proteins were induced with doxycycline (Dox) 16 h prior to the transfection with replicon RNA (n = 3-4). *(**c**)* Representative determination of Gaussia luciferase activity (mean of two technical replicates). **(*d*)** Quantification of luciferase activity 24 hours post transfection (hpt), normalized to the mean of the TAP-tag control cells (control). Error bars indicate SD (n = 3-4; 2 technical replicates measured per n; two-tailed Mann-Whitney test). *(**e**)* Whole cell lysates were analyzed for expression of ARTD10 (5H11) and ARTD12 (Sigma) by immunoblotting. HEK293 cells were transfected with *in vitro* transcribed RNA of the indicated CHIKV replicon variants (n = 3). *(**f**)* HEK293 Flp-In T-REx cells stably expressing the indicated TAP-tag proteins were transfected with plasmids encoding HA-tagged ARTD12 or ARTD12-HY. Twenty-four h later, the TAP-tag proteins were induced with Dox for 16 h and subsequently transfected with ^3^EGFP replicon RNA (n = 3). Gaussia luciferase activity was analyzed 9 and 24 hpt (left and right panel, respectively). Error bars indicate SD (n = 3; 2 technical replicates measured per n; two-tailed Mann-Whitney test). *(**g**)* Cells were treated as in panel c, infected with fully infectious virus expressing an EGFP reporter under the control of a subgenomic promotor with the indicated MOI and analyzed for GFP expression by flow cytometry 24 (left panel) and 48 h post infection (hpi) (right panel). All error bars indicate SD (n = 3, 3 technical replicates were measured per n; two-tailed Mann-Whitney test). (***P-values for panel a****: siControl vs. siARTD10 9 h P = 0.0003, 24 h P = 0.3282, 30 h P = 0.4418; siControl vs. siARTD12 9 h P = 0.0002, 24 h P = 0.0002, 30 h P = 0.0002; siControl vs. siARTD10/12 9 h P = 0.0003, 24 h P = 0.0002, 30 h P = 0.0002; siControl vs. siARTD8 9 h P = 0.2345, 24 h P = 0.0002, 30 h P = 0.0011; siControl vs. siARTD7 9 h P = 0.8785, 24 h P = 0.0281, 30 h P = 0.0002; **for panel d**:; control vs. ARTD10 P = < 0.0001; control vs. ARTD10-GW P = < 0.0001; control vs. ARTD12 P = 0.0002; control vs. ARTD12-HY P = 0.0559; ARTD10 vs. ARTD10-GW P = 0.0002; ARTD12 vs. ARTD12-HY P = 0.1320; **for panel f**: (left panel) control vs. ARTD12 P = 0.0022; control vs. ARTD12-HY P = 0.0022; control vs. ARTD10 P = 0.0022; control vs. ARTD10+ARTD12 P = 0.0022; control vs. ARTD10-GW P = 0.0152; ARTD12 vs. ARTD12-HY P = 0.0152; ARTD12 vs. ARTD10+ARTD12 P = 0.0022; ARTD10 vs. ARTD10+ARTD12 P = 0.3095; ARTD10 vs. ARTD10-GW P = 0.0022; (right panel) control vs. ARTD12 P = 0.0022; control vs. ARTD12-HY P = 0.1320; control vs. ARTD10 P = 0.0022; control vs. ARTD10+ARTD12 P = 0.0022; control vs. ARTD10-GW P = 0.0022; ARTD12 vs. ARTD12-HY P = 0.0022; ARTD12 vs. ARTD10+ARTD12 P = 0.0022; ARTD10 vs. ARTD10+ARTD12 P = 0.0022; ARTD10 vs. GW P = 0.0411**; for panel g**: (left panel, MOI 0.2) control vs. ARTD10 P = < 0.0001; control vs. ARTD10-GW P = 0.0012; control vs. ARTD12 P = 0.1615; control vs. ARTD12-HY P = 0.0188; ARTD10 vs. ARTD10-GW P = < 0.0001; ARTD12 vs. ARTD12-HY P = 0.1615; (MOI 0.6) control vs. ARTD10 P = < 0.0001; control vs. ARTD10-GW P = 0.0244; control vs. ARTD12 P = 0.0400; control vs. ARTD12-HY P = 0.0503; ARTD10 vs. ARTD10-GW P = < 0.0001; ARTD12 vs. ARTD12-HY P = 0.1135; (right panel, MOI 0.2) control vs. ARTD10 P = < 0.0001; control vs. ARTD10-GW P = 0.0008; control vs. ARTD12 P = 0.0028; control vs. ARTD12-HY P = 0.0188; ARTD10 vs. ARTD10-GW P = < 0.0001; ARTD12 vs. ARTD12-HY P = 0.0142; (MOI 0.6) control vs. ARTD10 P = < 0.0001; control vs. ARTD10-GW P = 0.0188; control vs. ARTD12 P = < 0.0001; control vs. ARTD12-HY P = 0.0503; ARTD10 vs. ARTD10-GW P = < 0.0001; ARTD12 vs. ARTD12-HY P = 0.2973)*.

To expand on these findings, we determined the effects of ARTD10 and ARTD12 on the replication cycle of infectious CHIKV, which expresses EGFP from an additional subgenomic promoter (Fig. 1g) [47]. Overexpression of ARTD10 and ARTD12 resulted in a significant decrease of EGFP expressing cells as measured by flow cytometry 24 and 48 hpi (Fig. 1g and Supplementary Fig. 2). In this set-up, the catalytically inactive mutant of ARTD12 restricted replication to the same extend as the wt protein, indicating that ARTD12 may have more than one mode of action. In contrast, ARTD10-GW enhanced viral replication, hinting at a dominant negative effect and thereby demonstrating dependency on catalytic activity of ARTD10 as a CHIKV restriction factor. Of note is also that the inhibitory effects of ARTD10 and ARTD12 are more pronounced at early time points (24 hpi) compared to later time points (48 hpi). Taken together our findings indicate, that MARylation driven by the IFNα responsive ARTD10 and ARTD12 restricted CHIKV replication.

### MAR interferes with polyprotein processing

To gain insight into a possible mechanism of how MARylation by ARTD10 and ARTD12 might interfere with viral replication, we determined the abundance of auto-proteolytically processed nsP3. Therefore, we made use of EGFP-encoding variants of the replicon (^2^EGFP and ^3^EGFP, in which EGFP is integrated after amino acids 466 or 383 in nsP2 or nsP3, respectively Fig. 1b and [48]), enabling us to visualize processed nsP2 or nsP3 proteins using a GFP-specific antibody (Fig. 2). HEK293 cells stably expressing ARTD10 or ARTD10-GW were employed. These cells were then transfected with or without ARTD12 expressing constructs and induced for ARTD10 expression prior to transfection with ^3^EGFP replicon RNA. The analysis of whole cell lysates showed a reduction in processed nsP3 in presence of either the enzymatically active ARTD12 or ARTD10. NsP3 was further reduced when both enzymes were expressed (Fig. 2a). These findings led us hypothesize that MARylation hampers polyprotein processing. Proper polyprotein processing is a prerequisite for RNA replication [43], consistent with impaired replication of a mutant replicon with an inactive protease (nsP2-C478A/S482A, referred to as CASA) (Fig. 2b,c) [44]. Similarly, a functionally active macrodomain is needed for replication as substitution of key amino acids in the macrodomain (D10A, V33E, for details concerning the replicon construct see Fig. 1b) interfered with replication (Fig. 2b,c) [6, 9]. To analyze how the lack of a functional macrodomain compromised replication, we determined the abundance of proteolytically processed nsP2 (Fig. 2d, for the specificity of the antibody see Supplementary Fig. 3a,b). As expected, nsP2 was detectable after transfection of the wildtype (wt) but not the CASA mutant replicon (Fig. 2d). Similarly, nsP2 was not detectable, when expressed from hydrolase deficient replicons (Fig. 2d), implying a defect in nsP2-mediated polyprotein processing in the absence of MAR hydrolase activity. This observation was corroborated with the ^3^EGFP and ^2^EGFP replicons and mutants thereof (Fig. 2e-g). Although replication of the EGFP-encoding variants was reduced compared to the wt replicon, it remained dependent on functional protease and MAR hydrolase activities (Fig. 2e,f). As for nsP2, GFP-tagged processed nsP3 was not properly generated from the hydrolase deficient replicons (Fig. 2g). However, compared to the CASA mutant, at least in the longer exposure, some signal for processed nsP3 was detectable, implicating that the loss of MAR hydrolase activity did not completely abolish polyprotein processing as in case of the CASA mutant (Fig. 2g). This is also reflected by the significantly higher replication of the hydrolase deficient mutants compared to the CASA mutant (Fig. 2c), although replication in general was strongly impaired.

**Figure 2:**
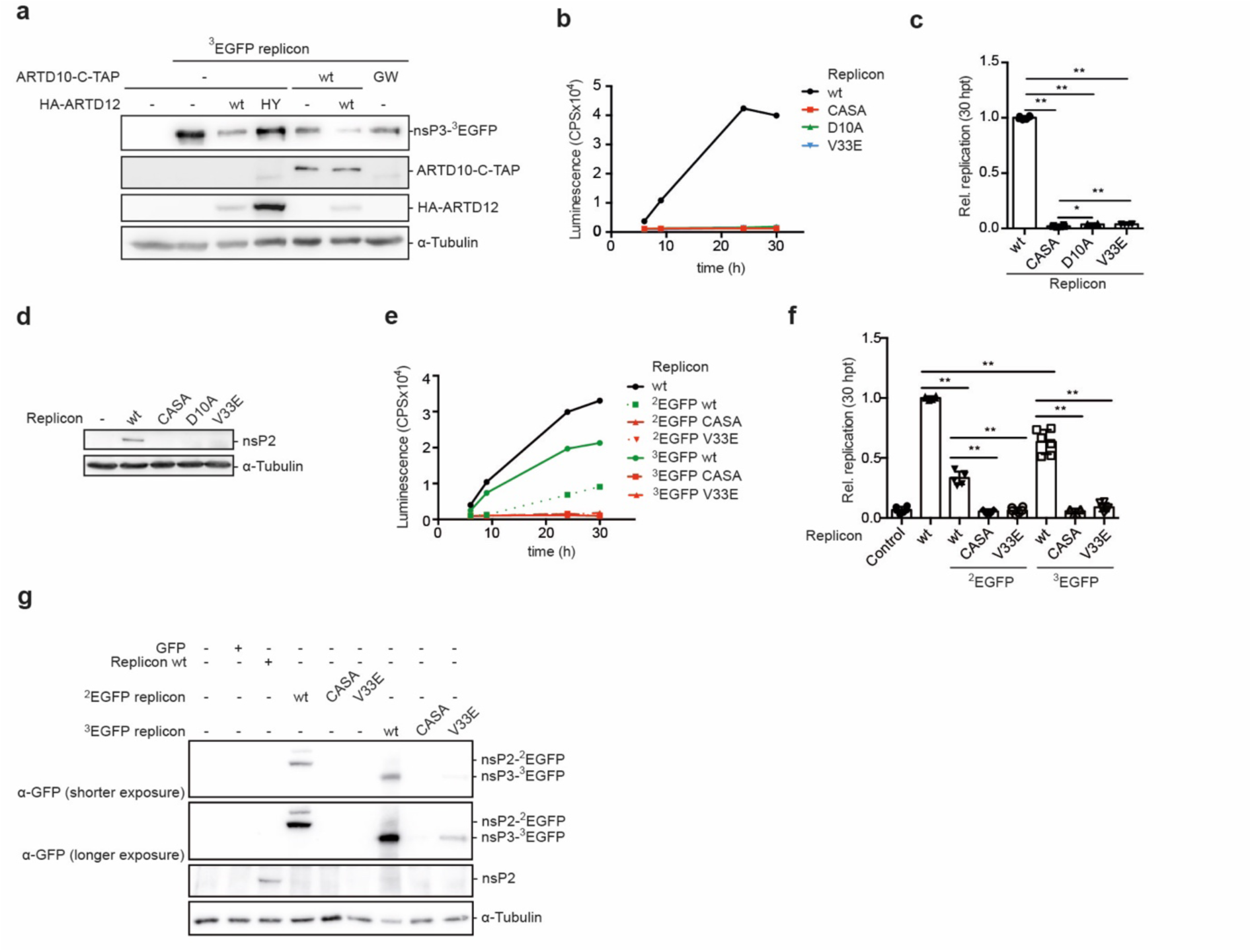
MARylation interferes with polyprotein processing. *(**a**)* HEK293 Flp-In T-REx cells stably expressing the indicated TAP-tag proteins were transfected with plasmids encoding HA-tagged ARTD12 or ARTD12-HY. Subsequently TAP-tag proteins were induced with Dox for 16 h. Twenty-four hpt the cells were transfected with ^3^EGFP replicon RNA. Whole cell lysates were analyzed for processed nsP3 (GFP) and expression of either ARTD10 (5H11) or ARTD12 (HA) by immunoblotting. *(**b**)* HEK293 cells were transfected with *in vitro* transcribed RNA of the wt replicon or mutants thereof. Representative measurement of Gaussia luciferase activity of the supernatants of the transfected cells in counts per second (CPS) at the indicated times (mean of two technical replicates). *(**c**)* Gaussia luciferase activity was analyzed as surrogate for CHIKV replication 30 hours post transfection (hpt) normalized to the mean of the wildtype (wt) for each experiment. All error bars indicate SD (n = 3; 2 technical replicates measured per n; two-tailed Mann-Whitney test). *(**d**)* Whole cell lysates of the cells analyzed in panel ***c*** were examined for processed CHIKV nsP2 by immunoblotting (n = 3). *(**e-g**)* HEK293 cells were transfected with the indicated CHIKV replicon variants or EGFP as control (n = 3). **(*e*)** Representative measurement of Gaussia luciferase activity of the supernatants of the transfected cells in counts per second (CPS) at the indicated times (mean of two technical replicates). *(**f**)* Quantification of several experiments as exemplified in panel e. Error bars indicate SD (n = 3; 2 technical replicates measured per n; two-tailed Mann-Whitney test). *(**g**)* Whole cell lysates were analyzed for processed nsP3 (GFP) and nsP2 (GFP) by immunoblotting. (***P-values for panel c****: wt vs. CASA P = 0.0022; wt vs. D10A P = 0.0022; wt vs. V33E P = 0.0022; CASA vs. D10A P = 0.0152; CASA vs. V33E P = 0.0043; D10A vs. V33E P = 0.9372; **for panel f**: wt vs. ^2^EGFP wt P = 0.0022; ^2^EGFP wt vs. ^2^EGFP CASA P = 0.0022; ^2^EGFP wt vs. ^2^EGFP V33E P = 0.0022; wt vs. ^3^EGFP wt P = 0.0022; ^3^EGFP wt vs. ^3^EGFP CASA P = 0.0022; ^3^EGFP wt vs. V33E 0.0022)*.

### Polyprotein synthesis itself is unaffected by replicon substitutions

To exclude that protein degradation is affected by MARylation, we determined replication and processed nsP2 or nsP3 in presence of either the proteasome inhibitor MG-132 or Bafilomycin A1, which inhibits fusion of autophagosomes with lysosomes (Fig. 3a,b). Inhibition of neither pathway was able to rescue replication and nsP processing (Fig. 3a,b), supporting our hypothesis that polyprotein processing is affected. To expand on this, we verified that the synthesis of the polyprotein was not influenced by the individual mutations. In the absence of suitable antibodies to detect the full-length polyprotein on Western blots, our own and commercially available failed to do so, we turned to complementation experiments (Fig. 3c-n, Supplementary Fig. 4). We found that the untagged CASA replicon with its functional macrodomain was capable of rescuing replication of the hydrolase deficient ^3^EGFP replicon and vice versa (Fig. 3c,d). Next, we analyzed GFP rather than luciferase expression by immunoblotting and flow cytometry to be able to distinguish between the hydrolase deficient and the protease deficient replicons. In line with the replication, an increase in processed nsP3 was detected (Fig. 3e). In addition, co-transfection of the CASA replicon resulted in a more than 50% rescue of GFP positive cells with increased signal intensity compared to the V33E mutant alone in flow cytometry analysis (Fig. 3f-h, Supplementary Fig. 5).

**Figure 3:**
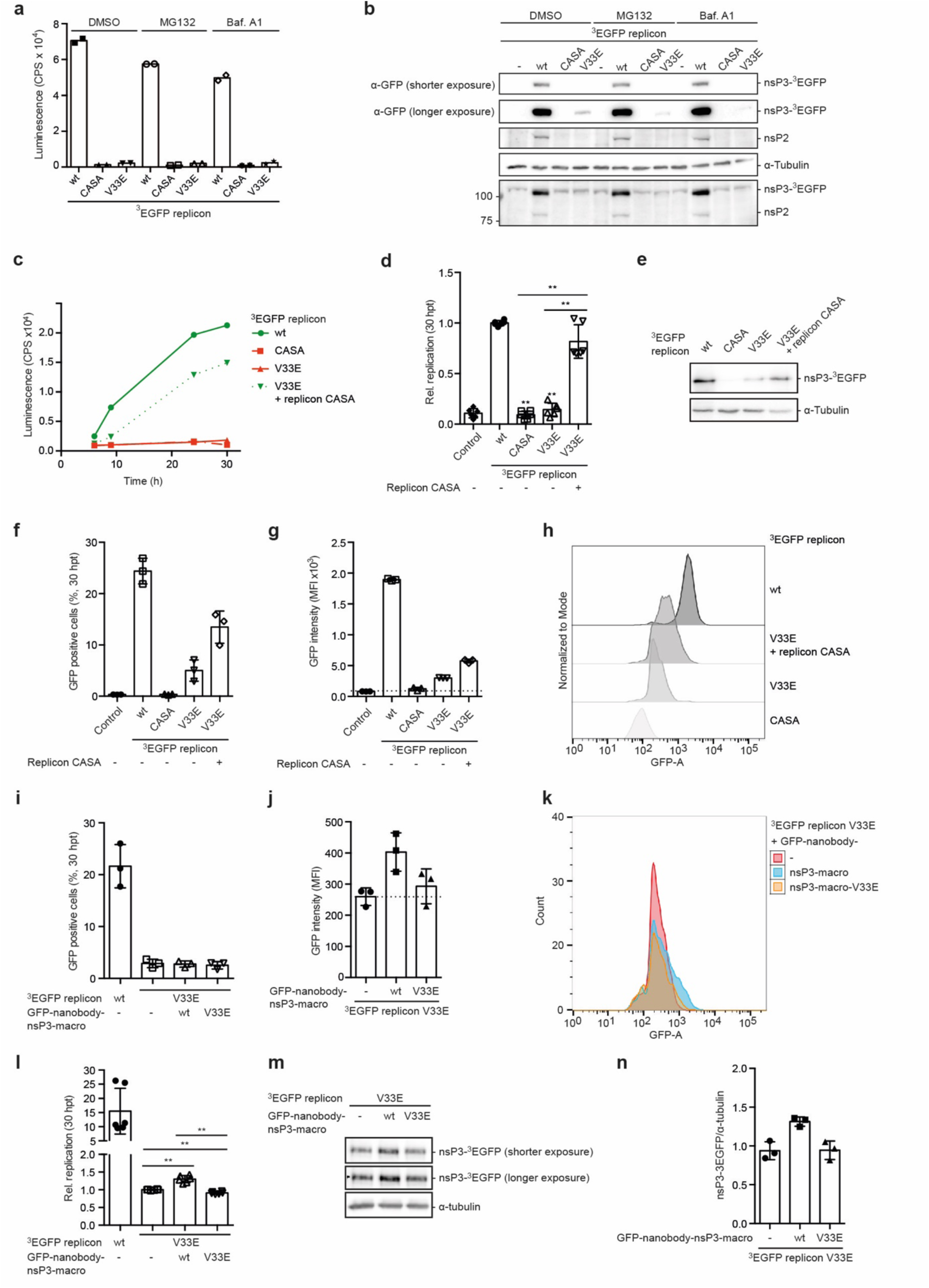
Polyprotein processing and viral replication requires both protease and MAR hydrolase activity. *(**a,b**)* HEK293 cells were transfected with *in vitro* transcribed ^3^EGFP wt and mutant replicons as indicated. Twenty-four hpt, the cells were treated with either vehicle, MG132 or Bafilomycin A1 (Baf.A1) for 6 h to inhibit proteasomal or lysosomal protein degradation, respectively. (**a**) Gaussia luciferase was measured in duplicates at 30 hpt. *(**b**)* Whole cell lysates were analyzed for processed nsP2 and nsP3 (GFP) by immunoblotting (n = 1). *(**c-h**)* HEK293 cells were (co-)transfected with RNA of the ^3^EGFP replicons and the replicon CASA mutant as indicated (n = 3). *(**c**)* Representative measurement of Gaussia luciferase activity of the supernatants of the transfected cells in counts per second (CPS) at the indicated times (mean of two technical replicates). *(**d**)* Gaussia luciferase activity normalized to the mean of the ^3^EGFP replicon wt for each experiment 30 hpt. Error bars indicate SD (n = 3; 2 technical replicates measured per n; two-tailed Mann-Whitney test). *(**e**)* Whole cell lysates were analyzed for processed nsP3 (GFP) by immunoblotting. *(**f,g**)* Flow cytometry was used to determine GFP positive cells (***f***) as well as their mean GFP fluorescence intensity (MFI) 30 hpt (***g***) (n = 3). *(**h**)* Representative visualization of the MFI of the GFP positive cells from panels ***f*** and ***g*** with the “Modal” option scaling all channels to a percentage of the maximum count. *(**i-n**)* HEK293 cells were transfected with plasmids encoding anti-GFP-nanobody-nsP3-macro wt or V33E mutant. Twenty-four h later cells were transfected with *in vitro* transcribed ^3^EGFP wt and V33E mutant replicons as indicated. *(**i,j**)* Flow cytometry was used to determine GFP positive cells as well as their MFI 30 hpt (n = 3). *(**k**)* Representative visualization of the GFP intensity of the GFP positive cells. *(**l**)* Gaussia luciferase activity normalized to the mean of the ^3^EGFP replicon V33E for each experiment 30 hpt. Error bars indicate SD (n = 3; 2 technical replicates measured per n). *(**m,n**)* Whole cell lysates were analyzed for processed nsP3 (GFP) by immunoblotting and the amount of nsP3-^3^EGFP was quantified by densitometry in relation to the loading control α-tubulin (n = 3). (***P-values for panel d****: wt vs. CASA P = 0.0022; wt vs. V33E P = 0.0022; wt vs. V33E+CASA P = 0.3095; CASA vs. V33E+CASA P = 0.0022; V33E vs. V33E+CASA P = 0.0022; **panel l**: V33E vs. V33E + anti-GFP-nanobody-nsP3-macro P = 0.0022; V33E vs. V33E + anti-GFP-nanobody-nsP3-macro-V33E P = 0.0022; anti-GFP-nanobody-nsP3-macro vs. anti-GFP-nanobody-nsP3-macro-V33E P = 0.0022)*.

Then we examined whether replication could also be rescued by co-expression of plasmid-encoded nsP2 or nsP3 and the respective replicon mutant RNA (CASA or V33E) (Supplementary Fig. 4). Co-expression of the nsP2-459-798 protease domain partially rescued replication of the CASA mutant suggesting that indeed polyprotein synthesis takes place (Supplementary Fig. 4a-c). In contrast, co-expression of nsP3 or the isolated nsP3 macrodomain was not sufficient to rescue replication of a hydrolase deficient replicon mutant (Supplementary Fig. 4d-f). To overcome possible differences in the subcellular localization of replication hubs and plasmid-expressed nsP3 macrodomain, we fused the macrodomain to an anti-GFP-nanobody to enhance targeting to sites of replication of the EGFP encoding replicons. We transfected HEK293 cells with plasmids coding for a GFP-nanobody-nsP3-macrodomain fusion protein prior to transfection with the ^3^EGFP-replicon RNA. We analyzed GFP by flow cytometry (Fig. 3i-k, Supplementary Fig. 5). Whereas the overall amount of GFP-positive cells when transfected with the V33E mutant was low compared to the wildtype and stayed unaffected by co-expression of GFP-nanobody-nsP3-macrodomain (Fig. 3i), we observed an increase in GFP intensity dependent on hydrolytic activity of the GFP-nanobody fusion protein (Fig. 3j,k, Supplementary Fig. 5). In line with this replication in presence of the wildtype GFP-nanobody-nsP3-macrodomain was slightly but significantly increased (Fig. 3l). Further, the analysis of nsP3 revealed an increase in its processed form (Fig. 3m,n), indicating that the co-expression of the GFP-nanobody-nsP3-macrodomain at least to some extend was able to rescue the CHIKV replicon lacking hydrolase activity.

Taken together, these data suggest strongly that polyprotein synthesis of the different mutated replicons occurs and further support the hypothesis that MARylation affects polyprotein processing.

### CHIKV nsP2 is a substrate for MARylation *in vitro* and in cells

Consequences of protein MARylation are poorly understood. Our previous studies indicated that ARTD10-dependent MARylation impairs the catalytic activity of the kinase GSK3β, which is antagonized by cellular MAR hydrolases [37, 49]. Furthermore, MARylation is reported to affect protein-protein interactions, mRNA stability and translation [1]. Following our hypothesis, we determined the consequences of MARylation on the protease activity of nsP2, which is responsible for polyprotein processing [43]. Therefore, we tested whether nsP2 serves as substrate for mono-ARTDs. His_6_-tagged fusion proteins of CHIKV nsP2 or nsP2-459-798, comprising the protease domain, were incubated with His_6_-tagged catalytic domains of ARTD7, ARTD8, ARTD10 and ARTD12 (Fig. 4a, Supplementary Fig. 6). The corresponding genes are IFNα responsive (Supplementary Fig. 1a and [6]). Moreover, we tested ARTD15, which is not regulated by IFNα (Supplementary Fig. 1a). For all ARTD catalytic domains auto-ADP-ribosylation was measurable in presence of ^32^P-NAD^+^ even though the signal intensities varied considerably between the different enzymes (Fig. 4a, Supplementary Fig. 6a, b) [14, 16]. Both, full length CHIKV nsP2 as well as the isolated protease domain were MARylated by the catalytic domains of IFN-regulated ARTDs but not of ARTD15 (Figure 4a, Supplementary Fig. 6a).

**Figure 4:**
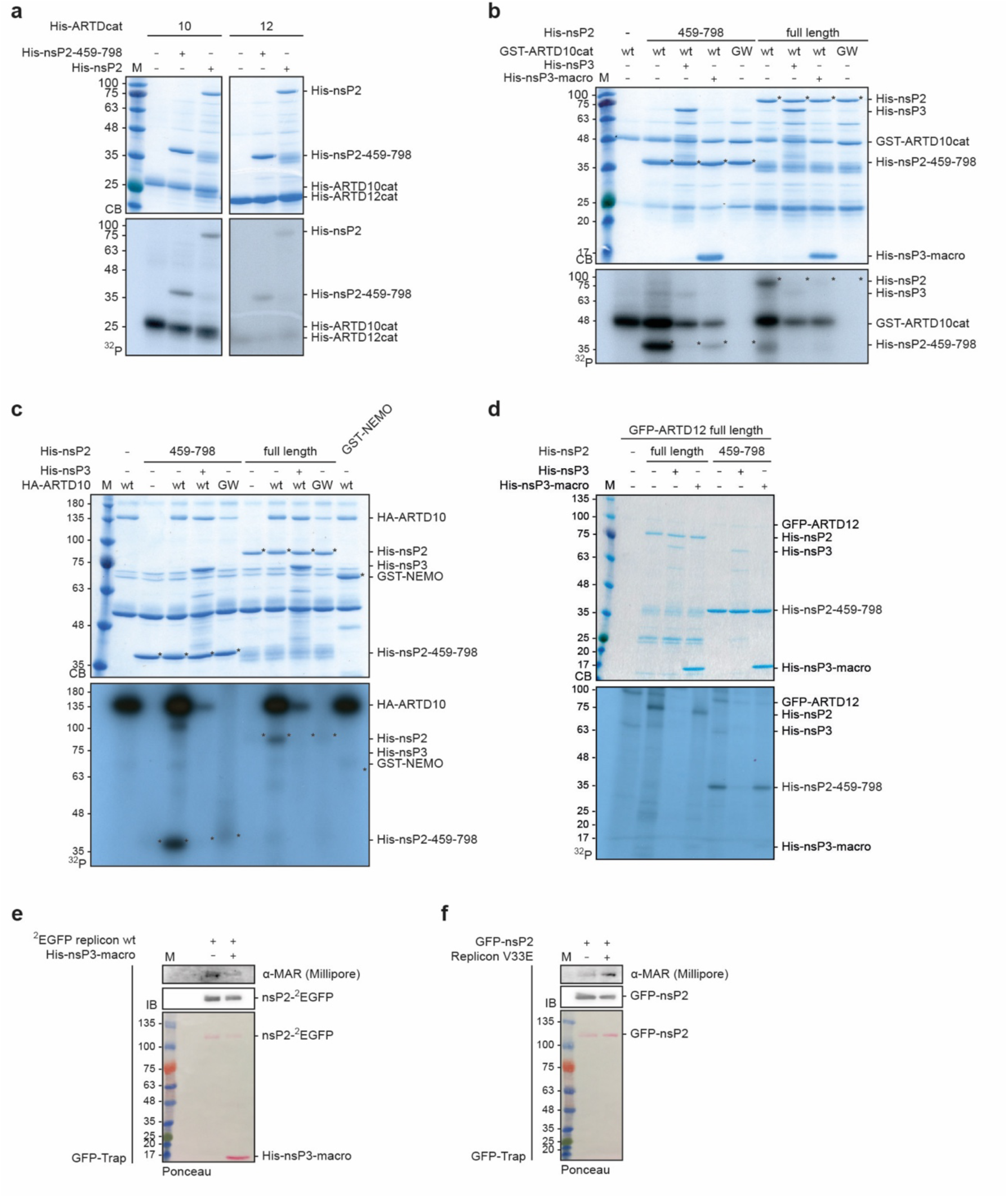
Nsp2 is MARylated by mono-ARTDs and de-MARylated by the nsP3 macrodomain. *(**a**)* Bacterially expressed and purified His_6_-tagged ARTD catalytic domains (cat) and His_6_-tagged CHIKV nsP2 or its protease domain (nsP2-459-798) were subjected to *in vitro* ADP-ribosylation assays using ^32^P-NAD^+^ for 30 min at 30°C. The reactions were subjected to SDS-PAGE and the proteins were stained using Coomassie blue (CB). The incorporated radioactive label was assessed by autoradiography (^32^P) (the gel with all ARTDs analyzed is shown in Fig. S4a) (n = 2). *(**b**)* Bacterially expressed and purified GST-ARTD10cat and His_6_-tagged CHIKV or nsP2-459-798 were MARylated as in panel a. The catalytically inactive ARTD10-G888W (GW) served as a negative control. The samples were co-incubated with His_6_-tagged nsP3 or nsP3-macro. The proteins were visualized using CB and by autoradiography (n = 2). *(**c**)* HEK293 cells were transfected with HA-tagged ARTD10 or ARTD10-GW, lysed and the HA fusion proteins immunoprecipitated with an HA-specific antibody. The immunoprecipitated proteins were subjected to a MARylation assay as described in panel a and b (n = 2). *(**d**)* As in panel c with immunoprecipitated GFP-ARTD12 (n = 2). *(**e**)* HEK293 cells were transfected with the ^2^EGFP replicon. The cells were lysed and the EGFP fusion proteins were immunoprecipitated with GFP-TRAP-MA beads 30 hpt. The immunoprecipitates were incubated with or without His_6_-tagged nsP3-macro for 30 min at 30°C. The proteins were analyzed by immunoblotting with a MAR-specific reagent (n = 1). *(**f**)* HEK293 cells were transfected first with plasmids encoding EGFP-nsP2, 24 h later with the V33E replicon, and 30 h later the cells were lysed. The EGFP fusion proteins were immunoprecipitated with GFP-TRAP-MA beads and analyzed by immunoblotting using a MAR-specific reagent (n = 1).

The presence of nsP3 in ADP-ribosylation reactions using the catalytic domains of either ARTD10 or ARTD12 reversed MARylation of nsP2 and the protease domain. Similarly, the isolated macrodomain also efficiently de-MARylated nsP2 (Fig. 4b, Supplementary Fig. 6b). We obtained similar results with nsP2 MARylated by full length ARTD10, while no MARylation was obtained with ARTD10-GW (Fig. 4c). In addition, nsP2 was reversibly MARylated by full length ARTD12 (Fig. 4d). To complement these *in vitro* findings, we measured MARylation of nsP2 expressed in HEK293 cells transfected with the ^2^EGFP replicon (Fig. 4e). The immunoprecipitated nsP2-^2^EGFP stained positively with a MAR binding reagent. In support, this staining was reduced upon incubation with the recombinant nsP3 macrodomain, providing evidence that this protein is MARylated in cells (Fig. 4e). Similarly, upon co-transfection of a plasmid expressing GFP-nsP2 and the V33E replicon, nsP2 stained positively for MAR (Fig. 4f). Taken together, we identified CHIKV nsP2 as a new substrate for MARylation *in vitro* and in cells in context of a viral infection.

### MARylation of nsP2 reversibly inhibits its proteolytic activity

Having identified nsP2 as a new substrate for MARylation, we aimed at determining the consequences of this modification on proteolytic activity. Therefore, we established a protease assay using an nsP3/nsP4 junctional peptide (DELRLDRAGG|YIFSS) fused to GST and EGFP (Fig. 5a) [50–52]. Accessibility between the two globular tags was achieved by including a polylinker C-terminally of the cleavage site. This artificial substrate was cleaved by the recombinant nsP2 protease with up to 90% processing observed after 120 min, but not by the CASA proteolytically inactive mutant (Fig. 5b). Of note is that neither the C-terminal EGFP fragment (fragment 1) nor the N-terminal GST fragment (fragment 2) were further cleaved, supporting the specificity of the protease activity. Both, the substrate and the two nsP2 variants were stable when analyzed individually over 120 min (Fig. 5b).

**Figure 5:**
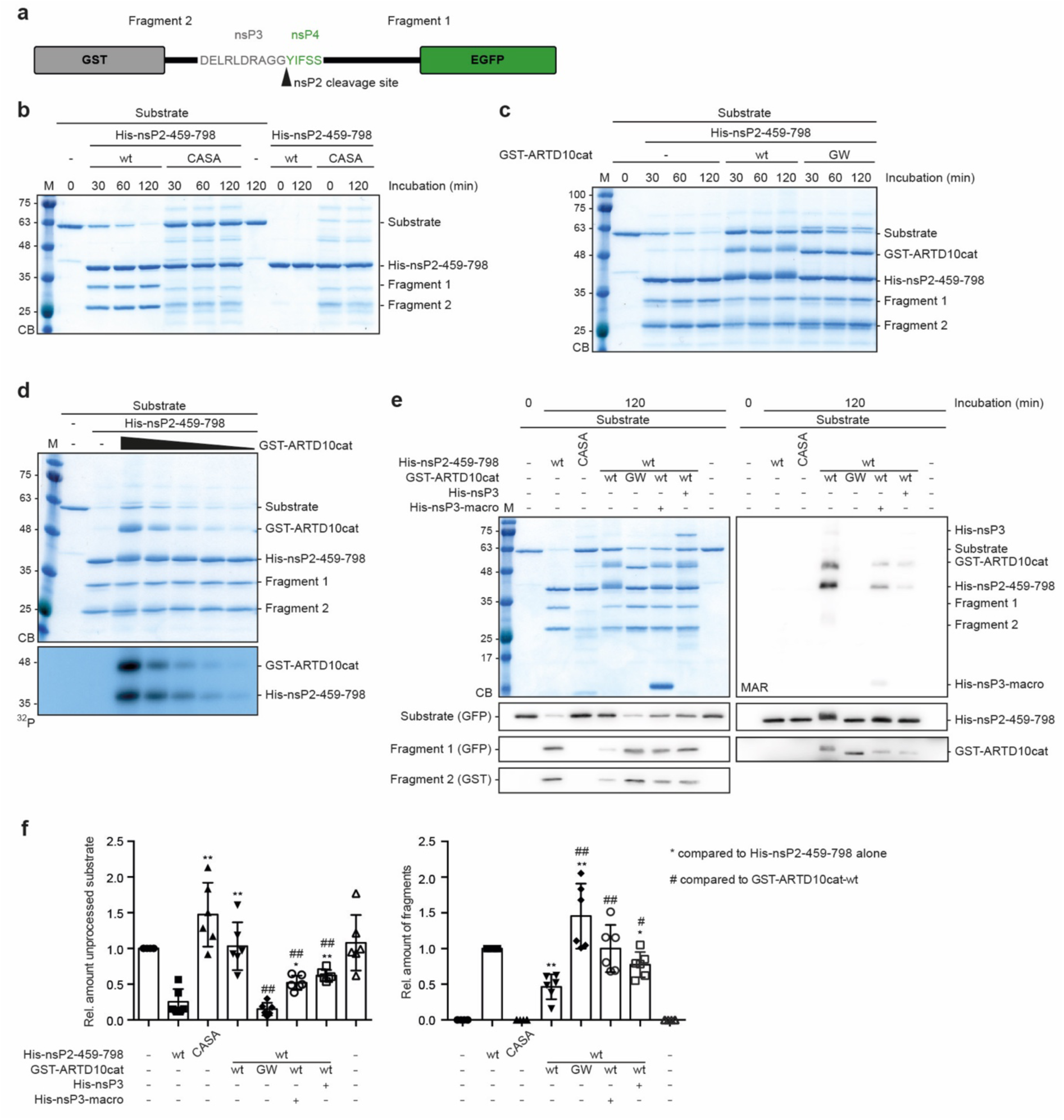
MARylation of nsP2 interferes with its protease activity. (**a**) Schematic representation of the synthetic nsP2 protease substrate with the nsP3/nsP4 cleavage site (long peptide described in [51]). (**b**) Bacterially expressed His_6_-tagged nsP2 protease domain (459-798) or the corresponding catalytically inactive CASA mutant were subjected to an *in vitro* protease assay with bacterially expressed and purified synthetic substrate at 30°C for the indicated times. The reaction products were subjected to SDS-PAGE and the proteins were stained with Coomassie blue (CB) (n = 2). (**c**) GST-ARTD10cat or the GW mutant and His_6_-tagged CHIKV nsP2-459-798 were incubated with NAD^+^ at 30°C for 30 min. Subsequently substrate was added and further incubated for the indicated times. Proteins were analyzed by SDS-PAGE and CB staining (n = 3). (**d**) nsP2-459-798 was incubated with increasing amounts of GST-ARTD10cat in presence of ^32^P-NAD^+^ at 30°C for 30 min. Then substrate was added for an additional 120 min. The proteins were detected by CB and by autoradiography (^32^P) (n = 1). (**e**) GST-ARTD10cat of the GW mutant, His_6_-tagged nsP2-459-798 or the CASA mutant, and His_6_-tagged nsP3 or nsP3-macro were incubated with NAD^+^ at 30°C for 30 min, as indicated. Subsequently, substrate was added and further incubated for 120 min. The proteins were analyzed by CB or by immunoblotting using antibodies specific for GST, EGFP, ARTD10 or nsP2 and by blotting with the MAR-specific reagent (n = 6). (**f**) Quantification of the experiments in panel e. The substrate (left panel) or the sum of the fragments 1 and 2 (right panel) were quantified by densitometry of the immunoblots. Error bars indicate SD (n = 6; two-tailed Mann-Whitney test). (***P-values for panel f****: (left panel) His-nsP2-459-798 alone wt vs. CASA P = 0.0022; His-nsP2-459-798 alone vs. GST-ARTD10cat-wt P = 0.0022; His-nsP2-459-798 alone vs. GST-ARTD10cat-GW P = 0.3939; His-nsP2-459-798 alone vs. His-nsP3-macro P = 0.0260; His-nsP2-459-798 alone vs. His-nsP3 P = 0.0043; GST-ARTD10cat-wt vs. His-nsP2-459-798-CASA P = 0.1320; GST-ARTD10cat-wt vs. GW P = 0.0022; GST-ARTD10cat-wt vs. His-nsP3-macro P = 0.0087; GST-ARTD10cat-wt vs. His-nsP3 P = 0.0087; (right panel) His-nsP2-459-798 alone vs. GST-ARTD10cat-wt P = 0.0022; His-nsP2-459-798 alone vs. GST-ARTD10cat-GW P = 0.0022; His-nsP2-459-798 alone vs. His-nsP3-macro P = >0.9999; His-nsP2-459-798 alone vs. His-nsP3 P = 0.0476; GST-ARTD10cat-wt vs. GW P = 0.0022; GST-ARTD10cat-wt vs. His-nsP3-macro P = 0.0022; GST-ARTD10cat-wt vs. His-nsP3 P = 0.0152)*.

To assess the role of MARylation, the nsP2 protease domain was modified by ARTD10cat. This prevented cleavage of the substrate, while ARTD10cat-GW had no effect (Fig. 5c). Successful MARylation of nsP2-459-798 is indicated by its mobility shift on SDS-PAGE (Fig. 5c). The inhibition of the protease function by ARTD10cat was dose dependent (Fig. 5d). To interrogate whether de-MARylation was sufficient to reactivate protease activity, we co-incubated the substrate with nsP2-459-798, ARTD10cat and nsP3 or nsP3-macro as indicated (Fig. 5e). De-MARylation, which was evident in the presence of hydrolase either by the reduced mobility shift or by staining with the MAR reagent, reactivated protease activity (Fig. 5e). The processing efficiency was quantified by measuring the intensity of unprocessed substrate and the two fragments by immunoblotting and densitometric analysis. This documented that MARylation by ARTD10cat efficiently repressed nsP2-459-798 protease activity, which was antagonized by nsP3 or the isolated macrodomain (Fig. 5e,f). Thus, these findings demonstrate that the nsP2 protease domain is reversibly inhibited by MARylation and support our initial hypothesis that MARylation inhibits polyprotein processing. We provide a mechanism how MARylation antagonizes and consequently how the nsP3 macrodomain contributes to CHIKV replication.

## Discussion

Taken together, we established that ARTD10 and ARTD12 interfere with CHIKV RNA replication and identified CHIKV nsP2 as target for MARylation by IFN-inducible mono-ARTDs. Mechanistically, our results provide evidence for inhibition of the nsP2 protease function, which is essential for viral replication, by ARTD10-dependent MARylation. This results in a defect in CHIKV polyprotein processing and consequently in reduced replication. This MARylation-dependent inhibition of the protease activity is antagonized by the macrodomain of nsP3. Accordingly, the lack of MAR hydrolase activity prevents polyprotein processing. Thus, our findings elucidate a mechanism for the importance of a functional macrodomain for CHIKV replication.

ARTDs have been linked previously to restriction of virus replication [22]. In this context the best studied ARTD family member is ARTD13 (ZAP, Zinc-finger antiviral protein), which is able to bind viral RNA, promoting its decay or interfering with its translation [53]. Further, ARTD13, which is catalytically inactive, contributes to the establishment of an antiviral immune response by crosstalk with the miRNA pathway, thereby stimulating expression of antiviral proteins, and by amplifying RIG-I signaling [53]. Antiviral activities have also been assigned to ARTD10, ARTD12 and ARTD14. Overexpression of these ARTDs was shown to interfere with Venezuelan equine encephalitis virus (VEEV) replication [7]. Additionally, ARTD12 was described to restrict Sindbis Virus (SINV) and CHIKV replication amongst other RNA viruses [7]. Further ARTD10, ARTD12 and ARTD14 downregulate cellular translation in cells infected with VEEV, which in the case of ARTD12 depends on catalytic activity [8]. ARTD12 has also been identified to restrict Zika virus (ZIKV) replication, which is mediated by depletion of the ZIKV non-structural proteins NS1 and NS3 [26]. Dependent on its catalytic activity, ARTD12 seems to promote poly-ADP-ribosylation (PARylation) of these two viral proteins. PAR chains in turn are argued to serve as scaffold to recruit E3 Ubiquitin-ligases that modify NS1 and NS3 by K48-linked polyubiquitination and thereby enable their proteasomal degradation [26]. Indeed, this concept has already been established for PARylation mediated by ARTD5 and 6 (tankyrase 1 and 2, respectively) [1]. More than 70 substrates have been identified to be regulated through PAR-mediated poly-ubiquitination [54]. The proposed mechanism is that ARTD12, as it is limited to MARylation, modifies NS1 and NS3, which serves as a seeding event for polymer forming ARTDs, possibly ARTD5 or 6 [26]. However, the effect of MARylation on CHIKV replication seems to be different as demonstrated by experiments with inhibitors of proteasomal and lysosomal pathways in this study (Fig. 3a,b). In addition to the ZIKV proteins, the nucleocapsid protein of Coronavirus (CoV) was recently suggested to be ADP-ribosylated during infection [55]. It will be interesting to identify the enzyme that catalyzes this modification and moreover to define the molecular consequences, for example on capsid formation.

In summary, these different reports suggest that ADP-ribosylation may interfere with multiple viral functions. This is consistent with the observation that at least four of the 12 catalytically active mono-ARTDs are induced by type I IFNs (Supplementary Fig. 1a) [1, 6, 11]. The fact that the viral macrodomains display MAR hydrolase activity and thereby are able to reverse MARylation of viral as well as cellular substrates [6, 9, 10, 39] further supports the concept of (mono-)ADP-ribosylation as a key player in host-virus interactions. Macrodomain mutations, which disable hydrolase activity, interfere with SINV replication in neurons and prevent neuropathy in mice [56], decrease pathogenicity and modulation of the host immune response of CoV [11, 39], and severely impair viral replication of Hepatitis E virus (HEV) [57]. Similarly, CHIKV replication is dependent on the MAR hydrolase activity of the macrodomain (Figs. 2 and 3, Supplementary Fig. 4) [9, 45]. As pointed out above, in most cases it remains to be defined what the relevant substrates of mono-ARTDs and viral macrodomains are and how MARylation affects substrate functions. Our study provides a mechanistic explanation for the inhibitory role of MARylation in CHIKV replication and establishes the concept that ARTD10-dependent MARylation of the viral protease inhibits its function, which is antagonized by the viral macrodomain. Together, this links reversible MARylation of a key viral protein to host-virus interaction.

## Material and Methods

### Cell lines and cell culture

HeLa, HEK293, HEK293 Flp-In T-REx-nsP3, -nsP3-macro, -ARTD10, -ARTD10-G888W[14], - ARTD12, and -ARTD12-H564Y cells were cultivated in DMEM supplemented with 10% heat-inactivated fetal calf serum (FCS) at 37°C in 5% CO_2_. All HEK293 Flp-In T-REx cell lines were additionally supplemented with 5 µg/mL Blasticidin S (Invivogen) and 200 µg/mL Hygromycin B (Invivogen) for selection during every second passage. After thawing cells were regularly tested for mycoplasma by first purifying genomic DNA with the peqGOLD tissue DNA Mini Kit (peqlab) according to the manufacturer’s instructions and then the PCR reaction was performed for detection of mycoplasma DNA with specific primers (GPO-1: 5’-ACTCCTACGGGAGGCAGCAGTA-3’, MGSO: 5’-TGCACCATCTGTCACTCTGTTAACCTC-3’).

Plasmid DNA transfection of cells was performed using the calcium phosphate precipitation technique. Cells were transfected 48 h after seeding and 24 h prior to transfection with *in vitro* transcribed replicon RNA.

Cells were transfected with *in vitro* transcribed replicon RNA using Lipofectamine 2000 (Thermo Fisher Scientific) according to the manufacturer’s instructions. In short, cells were seeded in 12 well plates. For transfection 3 µg of *in vitro* transcribed RNA were dissolved in 100 µl OptiMEM and 5µl of Lipofectamine 2000 were added, mixed, vortexed and incubated at room temperature for 5 min before adding dropwise to the cells. 100 µl of supernatant were collected 6, 9, 12, 24 and/or 30 hours post transfection (hpt) for analysis of Gaussia luciferase activity. 30 hpt transfection cells were lysed in RIPA buffer (10 mM Tris, pH 7.4; 150 mM NaCl; 1% NP-40; 1% DOC; 0.1% SDS; Protease inhibitor cocktail (PIC)), fractionated by SDS-PAGE and subjected to immunoblotting or used for flow cytometry analysis.

To mediate a knockdown of the gene of interest, HEK293 cells were transiently transfected with siGENOME SMARTpools (Dharmacon) directed against Non-Targeting Control #2 (D-001206-14), *ARTD7* (M-017186-00), *ARTD8* (M-023583-02), *ARTD10* (M-014997-03), and *ARTD12* (M-13740-01) using HiPerFect Transfection Reagent according to the manufacturer’s instructions for 72 h prior to transfection with *in vitro* transcribed replicon RNA (as described above). In short, immediately after seeding cells were transfected with a mixture of 55 µl of OptiMEM and 5 µl of HiPerFect Transfection reagent per ml of medium and a final siRNA concentration of 20 nM.

HEK293 Flp-In T-REx cells were transfected with pcDNA5/FRT/TO-nsP3, -nsP3-macro, -ARTD12 or the H564Y mutant and pOG44 (Invitrogen) using the calcium phosphate precipitation technique and selected by treating the cells with 5 µg/mL Blasticidin S (Invivogen) and 200 µg/mL Hygromycin B (Invivogen).

Sixteen h prior to transfection or infection HEK293 Flp-In T-REx cell lines were induced with 1 µg/ml doxycycline to induce the expression of stably integrated TAP-tagged constructs. Afterwards cells were transfected with *in vitro* transcribed replicon RNA as described above or infected with full length virus as described below (see ‘Virus infection and analysis of replication’).

Twenty-four hpt with *in vitro* transcribed replicon RNA cells were treated with vehicle (DMSO), 25 µM MG132 (Sigma) or 200 nM Bafilomycin A1 (Baf.A1) (Enzo Life Sciences) for 6 h. Subsequently, supernatants were collected and cells were lysed with RIPA buffer and subjected to SDS-PAGE and immunoblotting for analysis.

### Reagents and antibodies

The following reagents were used: β-NAD^+^ (Sigma), ^32^P-NAD^+^ (Perkin-Elmer), IFNα (Peprotech), propidium iodide solution (Sigma), Protease inhibitor cocktail (Sigma), Glutathione-sepharose (Sigma), TALON metal affinity resin (BD Bioscience), GFP-Trap magnetic agarose beads (Chromotek, gtma), anti-GFP (Rockland, mouse monoclonal 600-301-215M and goat polyclonal 600-101-215), anti-α-Tubulin (Sigma, T5168 and Santa Cruz, sc-23948), anti-MAR binding reagent (Millipore, MABE1076), anti-GST (clone 6G9), anti-ARTD10 (clone 5H11 [14]), anti-ARTD12 (Sigma, SAB2104087), anti-Actin (clone C4, BP Biomedicals), anti-HA (BioLegend, clone 16B12), goat-anti-rabbit-HRP (Jackson Immunoresearch, 111-035-144), goat-anti-mouse-HRP (Jackson Immunoresearch, 115-036-068), goat-anti-rat-HRP (Jackson Immunoresearch, 112-035-068), rabbit-anti-goat-HRP (Santa Cruz, sc-2768).

Rabbit polyclonal, purified CHIKV-nsP2-specific antibodies were generated by immunizing rabbits simultaneously with two peptides (aa570-584: CERKYPFTKGKWNINK, and aa740-755: CVLGRKFRSSRALKPP), both located in the C-terminal third of CHIKV nsP2 (performed by Eurogentec).

### Cloning and mutagenesis

The SP6-CHIKV-replicon-SG-GLuc (hereafter referred to as replicon wt) construct was obtained from B. Kümmerer [46]. EGFP insertions were created on the basis of Utt et al 2016 [48]. Linkers (5’-ACTAGTTCCGAGCTCGAG-3’) with restriction sites for *Spe*I and *Xho*I were introduced by PCR based mutagenesis using the Q5 mutagensis kit (NEB) after codon 466 of nsP2 (^2^EGFP) or after codon 383 of nsP3 (^3^EGFP). The sequence encoding EGFP was amplified from pEGFP-C1 flanked by a *Spe*I restriction site and a Gly-Gly linker at the 5’-end and a Gly-Gly and a *Xho*I restriction site at the 3’-end by PCR and inserted into the linkers by restriction digestion and ligation. Single site mutations (C478A/S482A (CASA) in nsP2 and D10A or V33E in nsP3) were introduced into the replicon variants by insertion of custom-made DNA gBlocks (IDT). These were integrated by restriction digestion with *Nde*I for nsP2 or *Cla*I (5’-end) and *BstAP*I (3’-end) for nsP3 and ligation.

GST-ARTD10cat constructs were described previously [14]. pDest17-ARTD10cat constructs were created from pDONRZeo-ARTD10cat [14] using the gateway cloning system. The cDNAs encoding the catalytic domains of ARTD7 (N459-A656), ARTD8 (K1600-K1800), ARTD12 (G480-S688) and ARTD15 (N459-A656) were generated from plasmids obtained from H. Schüler (Stockholm) and cloned into pDest17 using gateway cloning (Thermo Fisher Scientific). pGEX4T1-ARTD12cat (489-684) was created from pNIC-28-BsaI-ARTD12 (M1-Q701) plasmid that was obtained from O. Gileadi (Oxford) by gateway cloning. pDest17-nsP3, pDest17-nsP3-macro and pGEX4T1-NEMO were described previously [6, 58]. pDest17-nsP2 and pDest17-nsP2-459-798 were generated with the gateway cloning strategy using the SP6-CHIKV-replicon-SG-GLuc as a template.

The artificial protease substrate (pGEX4T1-nsP3/nsP4-site-polylinker-EGFP) was created based on the long nsP3/nsP4 site described in Rausalu et al. [51]. This sequence was ordered as oligos containing *EcoR*I (5’-end) and *BamH*I (3’-end) restriction sites mimicking overhangs (5’-aattcGACGAGTTAAGACTAGACAGGGCAGGTGGGTATATATTCTCGTCGgag-3’, 3’-gatcctcCGACGAGAATATATACCCACCTGCCCTGTCTAGTCTTAACTCGTCg-5’) that were annealed *in vitro*. The sequence encoding EGFP was isolated from pEGFP-N1 using *BamH*I and *Not*I restriction sites and EGFP as well as the annealed oligos were inserted into pGEX4T1 using *Eco*RI and *BamH*I restriction sites and ligation. Subsequently a polylinker was introduced into this construct for better accessibility of the protease substrate. Therefore, oligos containing this polylinker, the nsP3/nsP4 site and *EcoR*I (5’-end) and *Nco*I (3’-end) restriction site mimicking overhangs (5’-aattcGACGAGTTAAGACTAGACAGGGCAGGTGGGTATATATTCTCGTCGGAGGATCCACCGGTCGCCACCGGCTCTGCCGCTGCCACAAGAGGCTCTGCTGGAAGCGGCGGATCTGCCACAGGCTCTGGATCTGCAGCTGGCTCTGGCGACTCTGTGGCTGCCGGATCTGGCGGAGGAAGCGGCTCTAc-3’, 3’-catggTAGAGCCGCTTCCTCCGCCAGATCCGGCAGCCACAGAGTCGCCAGAGCCAGCTGCAGATCCAGAGCCTGTGGCAGATCCGCCGCTTCCAGCAGAGCCTCTTGTGGCAGCGGCAGAGCCGGTGGCGACCGGTGGATCCTCCGACGAGAATATATACCCACCTGCCCTGTCTAGTCTTAACTCGTCg-5’) were annealed *in vitro* and inserted into the vector using *EcoR*I and *Nco*I restriction sites and ligation.

For the anti-GFP-nanobody constructs a human optimized sequence was ordered as oligos containing *Age*I (5’-end) and *Xho*I(3’-end) restriction sites (5’-ACCGGTCGCCACCATGCAGGTGCAGTTGGTAGAGAGTGGGGGAGCACTTGTTCAACCTGGAGGAAGTCTGCGGCTGTCATGCGCCGCCTCAGGCTTCCCGGTGAACAGATATTCCATGCGCTGGTACCGGCAAGCACCTGGCAAGGAGAGAGAATGGGTTGCAGGAATGAGTTCCGCAGGAGACAGAAGCAGCTATGA GGATTCTGTGAAAGGAAGGTTCACTATTAGCCGGGACGATGCACGGAACACTGTGTATCTCCAGATGAATTCCCTGAAGCCGGAGGATACGGCTGTCTACTATTGTAATGTAAATGTTGGATTCGAGTACTGGGGTCAAGGAACGCAAGTGACAGTATCCAGCTCCGGACTCAGATCTCGAG-3’). This sequence was inserted into GW-pEGFP-nsP3-macro or GW-pEGFP-nsP3-macro-V33E using the *Age*I and *Xho*I restriction sites and ligation, replacing the EGFP.

pEVRFO-HA and the pEGFP-ARTD10 constructs were described previously [14, 59]. pHA-, pEGFP-C1- and pcDNA5/FRT/TO-C-TAP-ARTD12 were created from the pNIC-28-BsaI-ARTD12 (M1-Q701) plasmid that was obtained from O. Gileadi (Oxford) by gateway cloning. Constructs for expression of eukaryotic fusion proteins of nsP2, nsP2-459-798, nsP3 and nsP3-macro were cloned into pcDNA3-Flag, pHA, pEGFP-C1 or pcDNA5/FRT/TO-N-TAP with gateway cloning using the SP6-CHIKV-replicon-SG-GLuc as a template. Mutants (except for replicon mutants, see above) were generated using standard mutagenesis procedures (e.g. Q5 mutagenesis kit (NEB)) and confirmed by sequencing. pcDNA3-HA-ARTD1 was a kind gift from M. Hottiger (Zürich) and pCMV-HA-ARTD14 from Andreas Ladurner (München).

### *In vitro* transcription of replicon RNA

For *in vitro* transcription of replicon RNA, DNA plasmids encoding the respective replicon variants were first linearized with *Nde*I. Subsequently, linearized DNA was transcribed using the mMESSAGE mMACHINE™ SP6 Transcription Kit (Thermo Fisher Scientific) according to the manufacturer’s instructions. Cap-analog [m^7^G(5’)ppp(5’)G] and GTP were added to the reactions to obtain 5’-capped RNA. Afterwards template DNA was digested by addition of TURBO DNase and RNA was precipitated using the lithium chloride precipitation protocol. Finally, RNA was resuspended in elution buffer from the High Pure RNA isolation Kit (Roche).

Purity was controlled by agarose gel electrophoresis, concentration was measured using a NanoDrop™ 1000 (Thermo Fisher Scientific) and RNA was stored at −80°C until transfection.

### Purification of His_6_- and GST-tagged fusion proteins

His_6_- and GST-tagged fusion proteins were expressed in *E. coli* BL-21. The recombinant proteins were enriched and purified via affinity chromatography on either glutathione-sepharose for GST-fusion or TALON metal affinity resin for His_6_-fusion proteins according to standard protocols. Purification of His-nsP2-459-798, wt or inactive CASA mutant, took place without the addition of PIC to the lysis buffer.

### Replicon assays

*In vitro* transcribed replicon RNA was transfected into cells as described above (see ‘*In vitro* transcription of replicon RNA and Cell lines and cell culture’). 100 µl of supernatants were collected 6, 9, 12, 24 and/or 30 hpt for analysis of Gaussia luciferase activity. Cells that were not transfected with replicon RNA functioned as negative control. Gaussia luciferase is under the control of the subgenomic promoter replacing the structural proteins and secreted into the supernatant [46]. Determining the Gaussia luciferase in the supernatant can thus function as a surrogate for CHIKV replication. To analyze the luciferase activity, the BioLux® Gaussia Luciferase Assay Kit (NEB, discontinued) or the GAR-2B Gaussia Luciferase Assay (Targeting Systems) were used according to the manufacturer’s instructions following the “Stabilized Assay Protocol I”. In short, 5 ml of dilution buffer were mixed with 800 µl of stabilizer and 50 µl of 100x substrate and incubated protected from light for 25 min at room temperature. Afterwards 5 µl of supernatant per sample were pipetted into a 96-well plate (opaque, white) in duplicates and mixed with 50 µl of substrate solution and incubated for 35-40 sec.

Afterwards the counts per second (CPS) were measured with a VICTOR^2^ 1420 multilabel counter (Perkin Elmer) measuring luminescence without a filter over 10 sec. To determine relative replication, values were normalized to the mean value of the 2 technical replicates of the according sample and time per experiment.

### Quantitative real-time PCR

To determine ISGs among the mono-ARTDs, HeLa cells were stimulated with IFNα (180 U/mL). Total RNA was isolated using the High Pure RNA isolation Kit (Roche) according to the manufacturer’s protocol. Reverse transcription was performed with 1 µg of the isolated RNA using the QuantiTect Reverse Transcription Kit (Qiagen). mRNA expression levels of *ARTD3*, *ARTD7*, *ARTD12*, *ARTD14* and *ARTD15* were analyzed by quantitative real-time PCR (qRT-PCR) using QuantiTect Primer Assays (QIAGEN). In all settings the mRNA expression of the gene of interest was normalized to *GUS* (forward 5’-CTCATTTGGAATTTTGCCGATT-3’ and reverse 5’-CCGAGTGAAGATCCCCTTTTTA-3’; IDT).

### *In vitro* ADP-ribosylation assays

ADP-ribosylation assays were performed in 30 µl reaction buffer (50 mM Tris, pH 8.0, 2 mM TCEP, 4 mM MgCl_2_) with 50 µM β-NAD^+^ and 1 µCi ^32^P-NAD^+^. After 30 min incubation at 30°C the reactions were stopped by addition of SDS sample buffer. Samples were fractionated by SDS-PAGE and gels subsequently stained with Coomassie blue to visualize the proteins. For the detection of the incorporated radioactive label, dried gels were exposed to X-ray films.

### *In vitro* ADP-ribosylation assays with immunoprecipitated ARTD10 and ARTD12

HEK293 cells were seeded and after 48 h transfected with plasmids encoding HA-ARTD10 or the inactive GW mutant or with plasmids encoding GFP-ARTD12 using the calcium phosphate precipitation technique. 48 hpt cells were lysed in TAP lysis buffer (50 mM Tris, pH 7.5; 150 mM NaCl; 1 mM EDTA; 10% glycerol; 1% NP-40; 2 mM TCEP; PIC) and the lysates were centrifuged at 4°C for 30 min. HA-ARTD10 was immunoprecipitated with 1 μl of anti-HA (BioLegend) antibody and protein G beads and GFP-ARTD12 with 5 µl of GFP-Trap magnetic agarose beads (Chromotek) at 4 °C for 1 h. Afterwards the beads were washed in TAP lysis buffer and reaction buffer (50 mM Tris, pH 8.0, 2 mM TCEP, 4 mM MgCl_2_). ADP-ribosylation assays were carried out as described above (chapter ***In vitro* ADP-ribosylation assays**).

### *In vitro* protease assay

Bacterially expressed and purified His-nsP2-459-798, wt or inactive CASA mutant, were incubated with synthetic substrate in 15 µl of reaction buffer (50 mM Tris, pH 8.0, 2 mM TCEP, 4 mM MgCl_2_) for 30, 60 or 120 min at 30°C. As a negative control substrate as well as proteases were incubated alone in reaction buffer for 0 or 120 min at 30°C. The reactions were stopped by the addition of SDS sample buffer. Samples were fractionated by SDS-PAGE and gels subsequently stained with Coomassie blue to visualize the proteins.

### ADP-ribosylation assay with subsequent *in vitro* protease assay

ADP-ribosylation assays were performed in 30 µl reaction buffer (50 mM Tris, pH 8.0, 2 mM TCEP, 4 mM MgCl_2_) with 50 µM β-NAD^+^ for 30 min at 30°C. Subsequently synthetic substrate was added to the reactions and they were further incubated at 30°C for 30, 60 or 120 min at 30°C. As a negative control substrate was incubated alone in reaction buffer for 0 or 120 min at 30°C. The reactions were stopped by the addition of SDS sample buffer. Samples were fractionated by SDS-PAGE and gels subsequently stained with Coomassie blue or subjected to immunoblotting to visualize the proteins.

### Immunoprecipitation for detection of MARylation in cells

HEK293 cells were seeded in 10 cm plates and 48 h after seeding transfected with plasmid DNA coding for GFP-nsP2 using the calcium phosphate precipitation technique or not treated. 24 h after DNA transfection or 72 h after seeding cells were transfected with *in vitro* transcribed replicon RNA as described above (chapter **Cell lines and cell culture**) but scaled up 10x according to the amount of medium. 30 hpt cells were harvested in RIPA buffer (10 mM Tris, pH 7.4; 150 mM NaCl; 1% NP-40; 1% DOC; 0.1% SDS; PIC) and the lysates were centrifuged at 4°C for 30 min. GFP-nsP2 or nsP2-^2^EGFP translated from the replicon RNA were immunoprecipitated with 5 µl GFP-Trap magnetic agarose beads (Chromotek) at 4°C for 1 h. Afterwards beads were washed in RIPA buffer and reaction buffer (50 mM Tris, pH 8.0, 2 mM TCEP, 4 mM MgCl_2_). Subsequent hydrolase assays were carried out with bacterially expressed and purified His-nsP3-macro in 10 µl reaction buffer for 30 min at 30°C. The reactions were stopped by the addition of SDS sample buffer. Samples were fractionated by SDS-PAGE and subjected to immunoblotting to visualize MARylation using the MAR reagent (Millipore) and the total proteins.

### Flow cytometry analysis

Thirty hpt with *in vitro* transcribed RNA or a plasmid encoding EGFP, cells were washed once and resuspended in 500 µl PBS containing 2% heat-inactivated FCS. For the propidium iodide (PI) single stain control, cells were then fixed and permeabilized in 80% ethanol for 30 min at −20°C and afterwards were washed twice and resuspended in 500 µl PBS containing 2% heat-inactivated FCS. All other samples were not fixed or permeabilized. Subsequently 50 µg/ml of PI solution (Sigma) were added to all samples and incubated in the dark for 20 min. A BD FACSCanto II (BD Bioscience) was used for the flow cytometry analysis of the samples. 100,000 events were counted per sample per experiment. Evaluation of the experiments was performed with the FlowJo software (BD Bioscience).

### Virus infection and analysis of replication

Infectious CHIKV, strain LR2006-OPY, was produced by *in vitro* transcription of the linearized full-length viral genome including an EGFP under a second subgenomic promotor [47] and subsequent electroporation of the RNA in BHK-21 cells. The virus was passaged once in BHK-21 cells. Infections were performed under BSL-3 conditions using MOI determined by titration on HEK293T cells. Cells were fixed in 4% PFA and infection efficiency was measured as the proportion of EGFP-positive cells 24 and 48 hours post infection by flow cytometry using a BD FACSLyric instrument (BD Bioscience). Evaluation of the experiments was performed with the FACSSuite v1.2.1.5657 software (BD Bioscience).

### Quantification of immunoblots and statistical analysis

Immunoblots were quantified using the Image J software (NIH, Bethesda, USA). The significance was analyzed using a nonparametric, two-tailed Mann-Whitney test with the GraphPad Prism software, since the number of values was too low to reliably test for Gaussian distribution of the data.

## Author contributions

S.K. performed the majority of the experiments; F.P. and C.G. performed the CHIKV full virus experiments; L.E. identified the IFNα-inducible mono-ARTDs by RT-qPCR; M.V. and M.B. purified recombinant proteins and B.Li. analyzed stable cell lines. S.K., P.V. and B.L. designed the experiments and wrote the paper. All authors read and approved the final manuscript.

## Acknowledgements

We thank A. Golzmann for expert technical assistance and B. Coutard, O. Gileadi, P.O. Hassa, M.O. Hottiger, B.M. Kümmerer and H. Schüler for providing plasmids. We thank H. Kleine for catalytically inactive ARTD12-expressing clones and A. Forst for generation of the stably expressing ARTD12 (wt/H564Y) Flp-In T-REx HEK293 cells. Further we thank F. Peisker and A. Bochyńska for support with the analysis of the flow cytometry data. This work was supported by the START program of the Medical School of the RWTH Aachen University (117/15) to P.V. and by the Deutsche Forschungsgemeinschaft DFG (VE1093/1-1, LU466/16-2) to P.V and B.L., respectively. C.G. is supported by a DFG grant within German-African Cooperation Projects in Infectiology (GO2153/3-1), by the Impulse and Networking Fund of the Helmholtz Association through the HGF-EU partnering grant PIE-008and by funding of Berlin Institute of Health (BIH).

