## supplementary files for "Mono-ADP-ribosylation by ARTD10 restricts Chikungunya virus replication by interfering with the proteolytic activity of nsP2"

*Patricia Verheugd

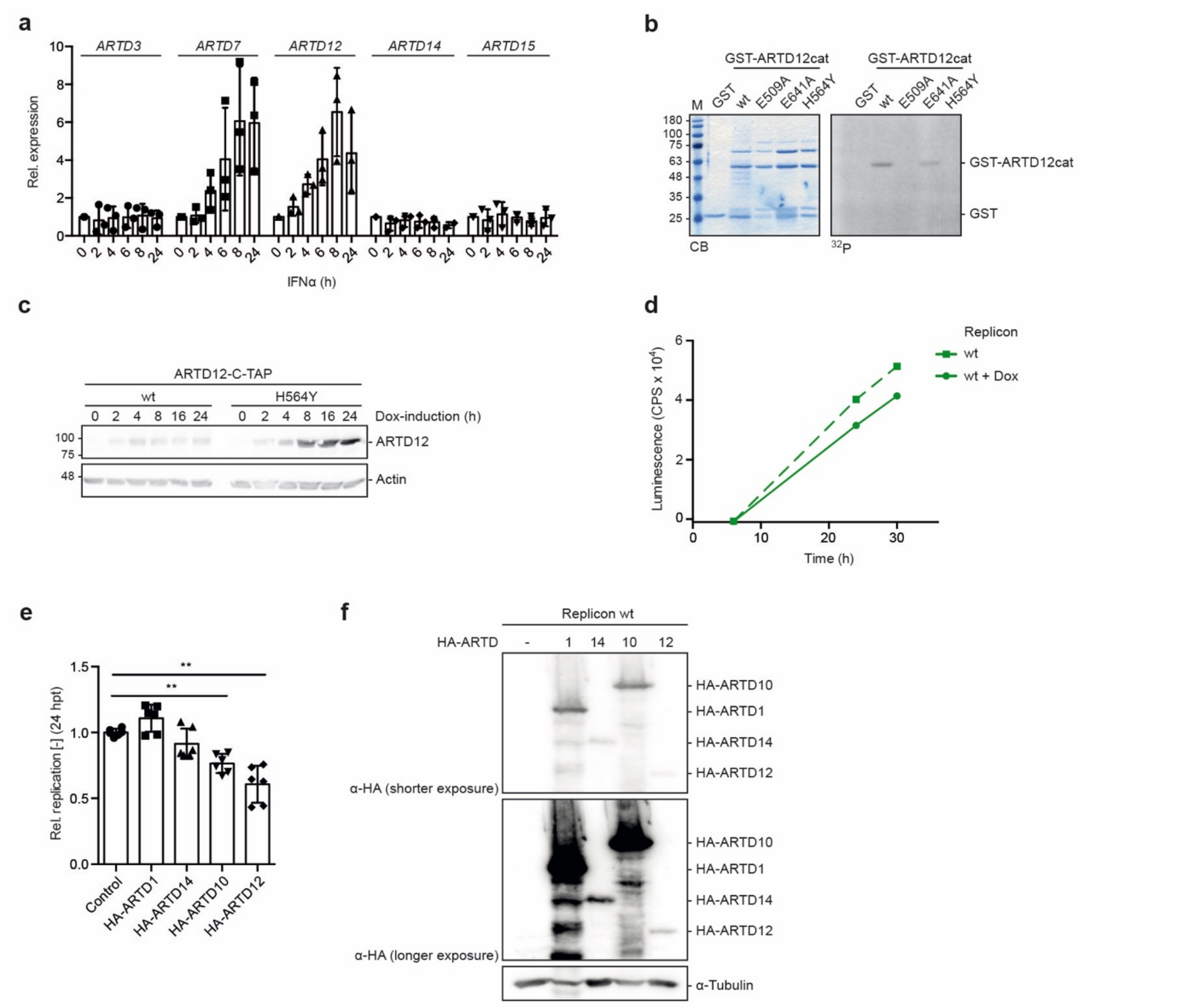


***Supplementary Figure 1 (supporting Figure 1a,c-g): (a)*** HeLa cells were stimulated with IFNα for the indicated times. The expression of *ARTD3*, *ARTD7*, *ARTD12*, *ARTD14* and *ARTD15* mRNA was analyzed using RT-qPCR and normalized to the unstimulated control. Error bars indicate SD (n = 3, 2 technical replicates were measured per n). ***(b)*** Bacterially expressed and purified ARTD12 or the indicated mutants were subjected to *in vitro* ADP-ribosylation assays in the presence of $^{32}$P-NAD$^{+}$ at 30°C for 30 min. The reactions were subjected to SDS-PAGE and the proteins were stained using CB and by autoradiography ($^{32}$P) (n = 1). ***(c)*** HEK293 Flp-In T-REx cells stably expressing ARTD12 or ARTD12-H564Y with a C-terminal TAP-tag were induced with doxycycline (Dox) for the indicated times. Proteins were analyzed with an ARTD12-specific antibody (n = 2). ***(d)*** Representative analysis of Gaussia luciferase activity in N-TAP HEK293 Flp-In T-REx cells treated with or without Dox (mean of two technical replicates). ***(e,f)*** HEK293 cells were transfected with plasmids encoding the indicated HA-tagged ARTDs and 24 h later with replicon RNA (n = 3). ***(e)*** Gaussia luciferase was determined 30 hpt, normalized to the control. Error bars indicate SD (n = 3; 2 technical replicates measured per n; two-tailed Mann-Whitney test). ***(f)*** Whole cell lysates were analyzed for expression of the HA fusion proteins by immunoblotting with an HA specific antibody 30 hpt.

(***P-values for panel e****: control vs. HA-ARTD1 P = 0.1797; control vs. HA-ARTD14 P = 0.3939; control vs. HA-ARTD10 P = 0.0022; control vs. ARTD12 P = 0.0022).*

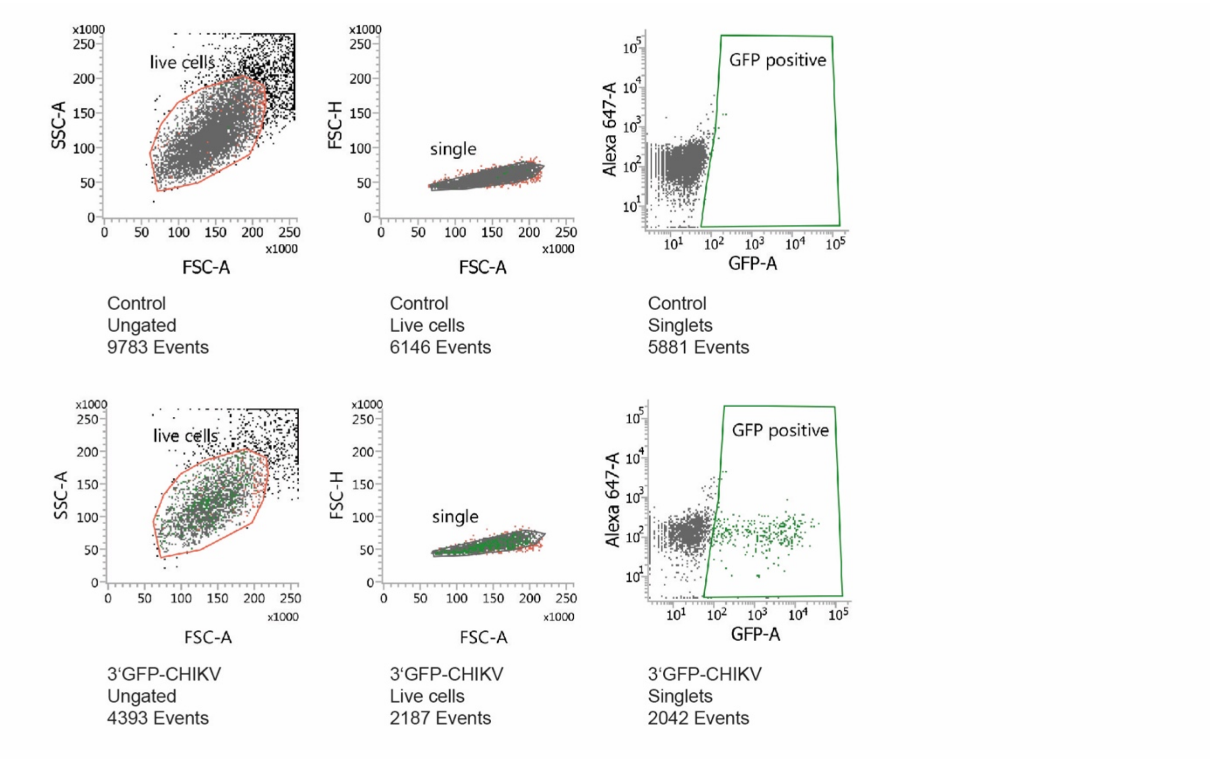


***Supplementary Figure 2 (supporting Figure 1g):*** HEK293 Flp-In T-Rex control cells or cells infected with fully infectious virus were analyzed by flow cytometry. Depicted are representative gating procedures used to analyze all samples. First the population was gated in SSC-A and FSC-A. Within this gate single cells were gated in the FSC-H and FSC-A.

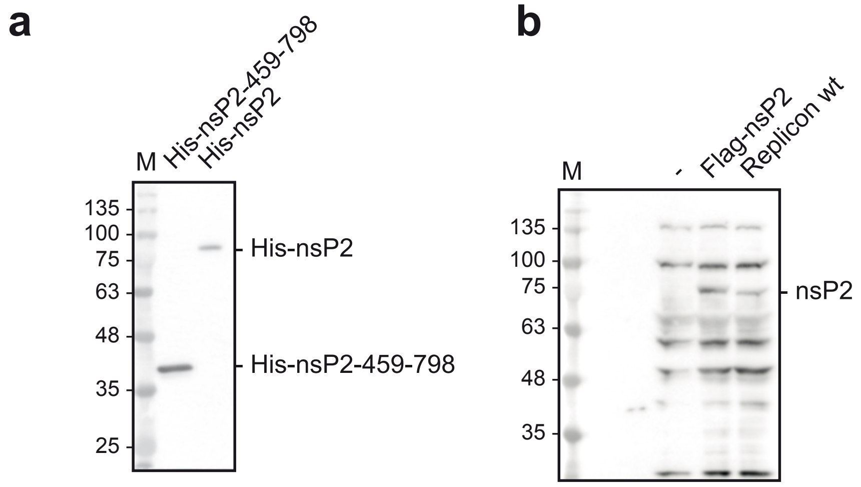


***Supplementary Figure 3 (supporting Figure 2d,g and 3b):*** Antibody evaluation of the custom-designed CHIKV-nsP2 antibody from Eurogentec. ***(a)*** Bacterially expressed and purified $His_6$-tagged fusions of either the isolated protease domain (His-nsP2-459-798) or the full length nsP2 were separated via SDS-PAGE and immunoblotted for testing the CHIKV nsP2 antibody. ***(b)*** HEK293 cells were transfected with either a plasmid encoding FLAG-nsP2 or *in vitro* transcribed wt replicon RNA. Whole cell lysates from transfected cells or control cells were analyzed via immunoblotting to evaluate CHIKV nsP2 protein in these lysates using the CHIKV-nsP2 antibody.

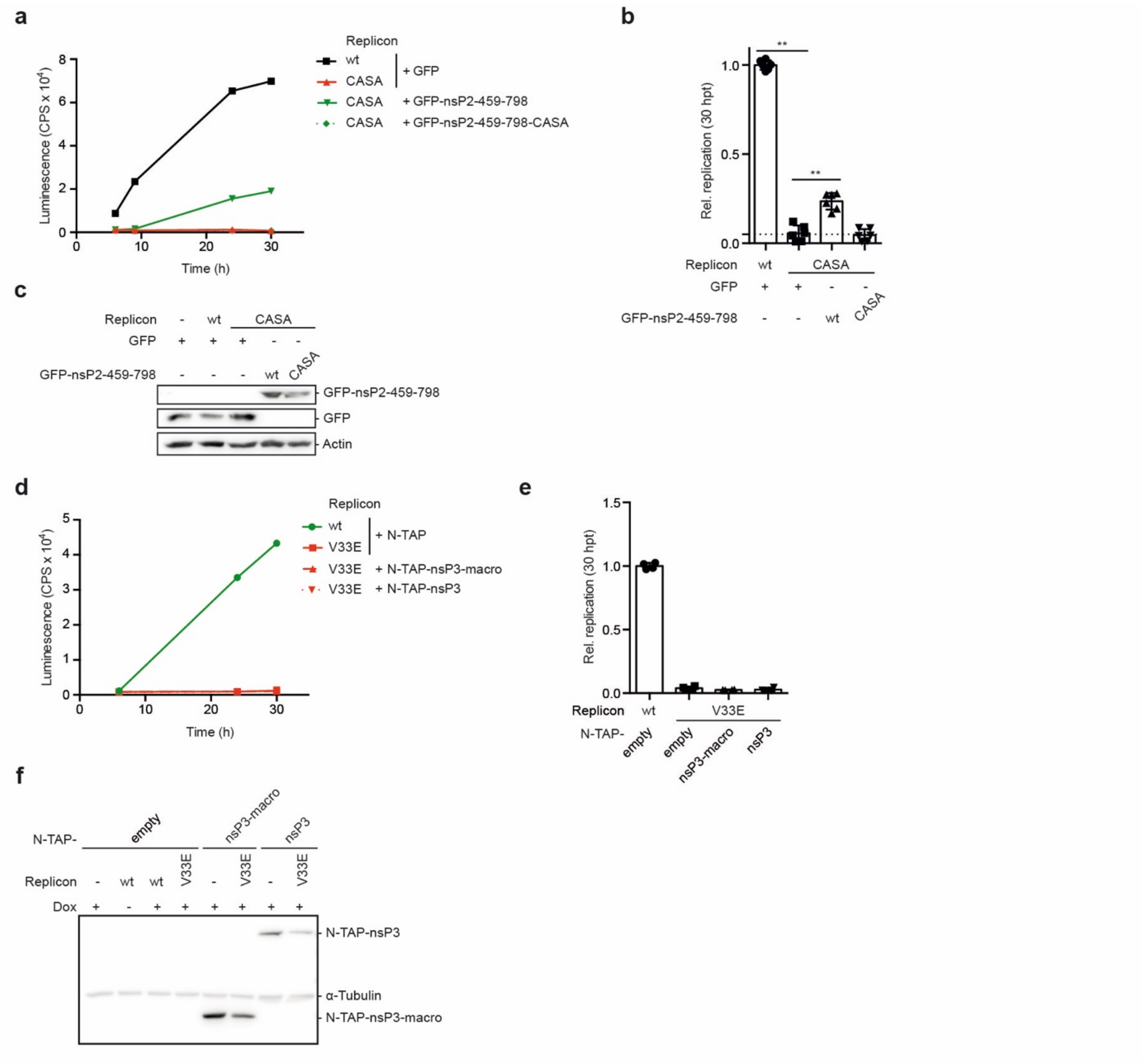


***Supplementary Figure 4 (supporting Figure 3):*** ***(a-c)*** HEK293 cells were co-transfected with *in vitro* transcribed RNA of the indicated replicon variants and plasmids encoding either EGFP or EGFP-nsP2-459-798 fusion proteins, with wt or CASA mutant protease domain, as indicated (n = 3). ***(a)*** Representative measurement of Gaussia luciferase activity (mean of two technical replicates). ***(b)*** Gaussia luciferase was determined 30 hpt, normalized to the wt replicon for each experiment. Error bars indicate SD (n = 3; 2 technical replicates measured per n; two-tailed Mann-Whitney test). ***(c)*** Whole cell lysates were analyzed for protein expression of the EGFP fusion proteins by immunoblotting. ***(d-f)*** HEK293 Flp-In T-REx cells stably expressing N-TAP, N-TAP-nsP3 or N-TAP-nsP3-macro were induced with doxycycline (Dox) 16 h prior to transfection with the indicated replicons (n = 2). ***(d)*** Representative determination of Gaussia luciferase activity from stable HEK293 cells upon induction of the indicated N-TAP proteins transfected with the indicated replicon (mean of two technical replicates). ***(e)*** Summary of luciferase activity of experiments as in panel g, error bars indicate SD (n = 2; 2 technical replicates measured per n). ***(f)*** Whole cell lysates were analyzed for expression of the stably expressing N-TAP fusion proteins by immunoblotting with an anti-rabbit secondary antibody. (***P-values for panel b****: wt + GFP vs. CASA + GFP P = 0.0022; CASA + GFP vs. CASA + GFP-nsP2-459-798 wt P = 0.0022; CASA + GFP vs. CASA + GFP-nsP2-459-798 CASA P = 0.5887).*

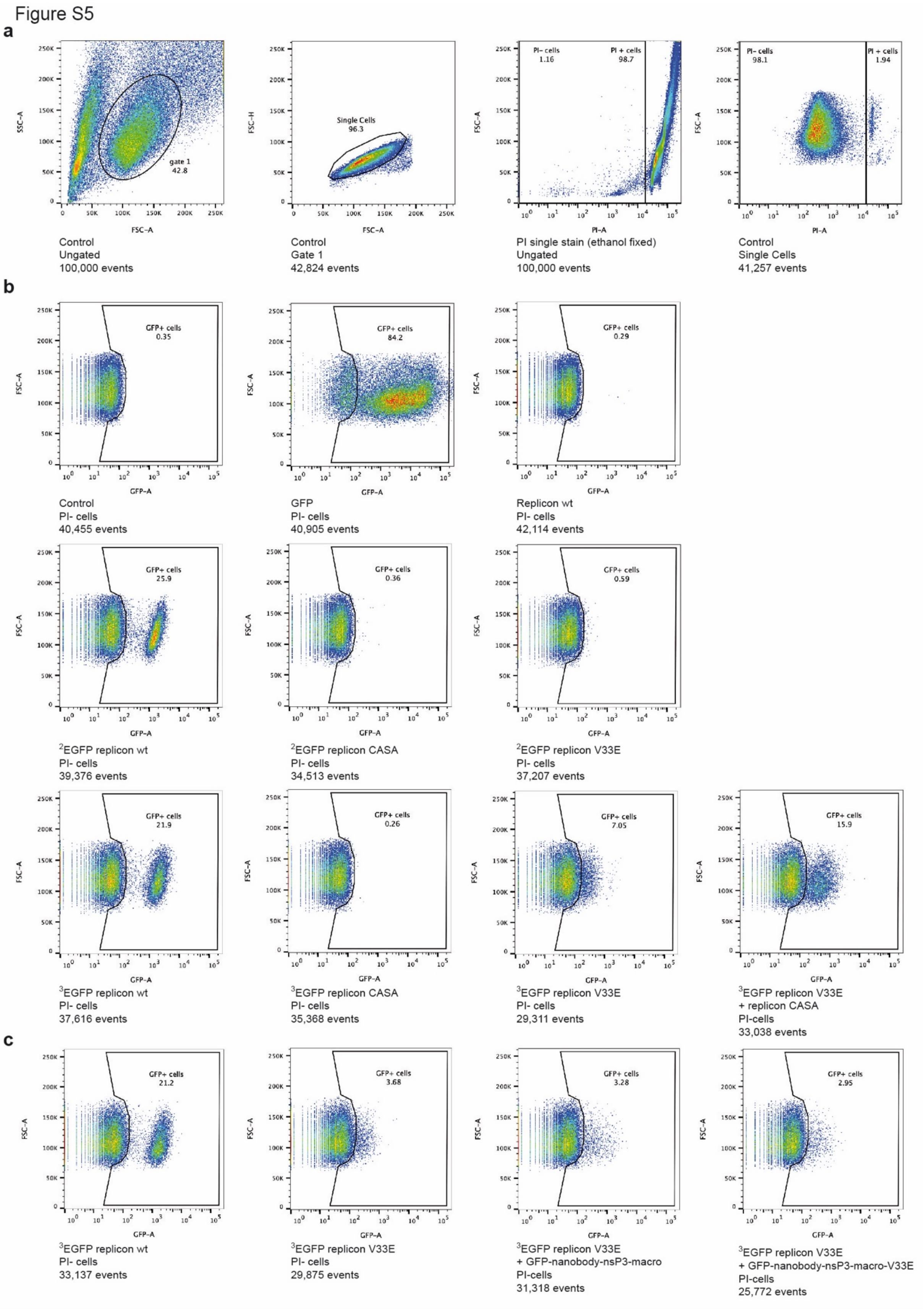


***Supplementary Figure 5 (supporting Figure 3f-k): (a,b)*** HEK293 cells were transfected with the indicated *in vitro* transcribed replicon RNA, a plasmid encoding EGFP as a control (Fig. 3f-h). Cells were analyzed by flow cytometry at 30 hpt and 100,000 events counted per experiment (n = 3). ***(a, c)*** HEK293 cells were transfected with the indicated GFP-nanobodies prior to transfection with *in vitro* transcribed RNA (Fig. 3i-k). Cells were analyzed by flow cytometry at 30 hpt and 100,000 events counted per experiment (n = 3). ***(a)*** Depicted is the representative gating procedure for control cells. The same gates were applied for all samples. First the population was gated in SSC-A and FSC-A. Within this gate single cells were gated in the FSC-H and FSC-A. To gate for the propidium iodide (PI) negative single cells, control cells were fixed and permeabilized with ethanol prior to PI staining. The according gates were applied for unfixed cells as well. ***(b,c)*** Depicted are representative gatings for GFP positive cells. Within the population of PI negative, single cells, a gate for GFP positive cells was set according to untransfected and EGFP transfected control cells. The gate was applied for all CHIKV replicon transfected cells and the percentage of GFP positive cells within the population is depicted.

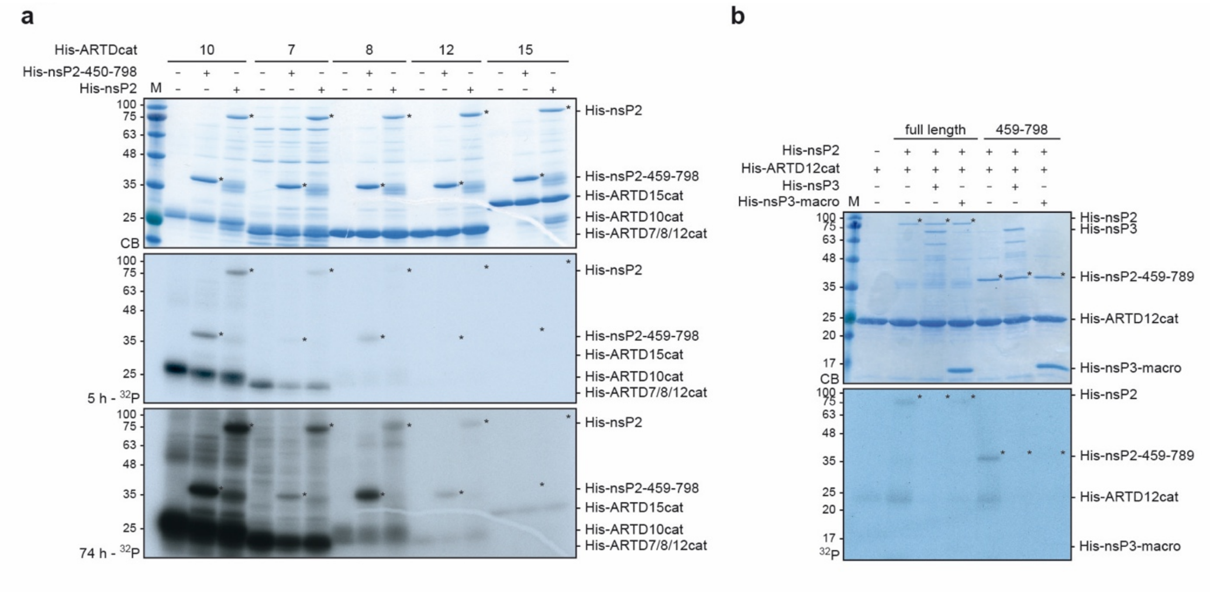


***Supplementary Figure 6 (supporting Figure 4):*** ***(a)*** Bacterially expressed and purified His$_6$-tagged ARTDcat and His$_6$-tagged CHIKV nsP2 or nsP2-459-798 were used in *in vitro* ADP-ribosylation assays with $^{32}$P-NAD$^+$ for 30 min at 30°C. The proteins were subjected to SDS-PAGE and stained using Coomassie blue (CB). The incorporated radioactive label was assessed by autoradiography ($^{32}$P) (n = 2). ***(b)*** Bacterially expressed and purified His$_6$-tagged ARTD12cat and His$_6$-tagged CHIKV nsP2 or nsP2-459-798 were used and analyzed as in panel a. In addition, His$_6$-tagged nsP3 or nsP3-macro were co-incubated (n = 2).
